# Leaps and bounds: geographical and ecological distance constrained the colonisation of the Afrotemperate by *Erica*

**DOI:** 10.1101/290791

**Authors:** M.D. Pirie, M. Kandziora, N.M. Nürk, N.C. Le Maitre, A. Mugrabi de Kuppler, B. Gehrke, E.G.H. Oliver, D.U. Bellstedt

## Abstract

The coincidence of long distance dispersal and biome shift is assumed to be the result of a multifaceted interplay between geographical distance and ecological suitability of source and sink areas. Here, we test the influence of these factors on the dispersal history of the flowering plant genus *Erica* (Ericaceae) across the Afrotemperate. We quantify similarity of *Erica* climate niches per biogeographic area using direct observations of species, and test various colonisation scenarios while estimating ancestral areas for the *Erica* clade using parametric biogeographic model testing. We infer that the overall dispersal history of *Erica* across the Afrotemperate is the result of infrequent colonisation limited by geographic proximity and niche similarity. However, the Drakensberg Mountains represent a colonisation sink, rather than acting as a “stepping stone” between more distant and ecologically dissimilar Cape and Tropical African regions. Strikingly, the most dramatic examples of species radiations in *Erica* were the result of single unique dispersals over longer distances between ecologically dissimilar areas, contradicting the rule of phylogenetic biome conservatism. These results highlight the importance of rare biome shifts, in which a unique dispersal event fuels evolutionary radiation.

This article has been peer-reviewed and recommended by: *Peer Community in Evolutionary Biology* (DOI: 10.24072/pci.evolbiol.100065)

## Introduction

The current day distributions of many plant groups are the result of long distance dispersal (LDD) (Muñoz et al., 2004; Alsos et al., 2007; Kadereit & Baldwin, 2012; Ruhfel et al., 2016; Jordano, 2017). Such events are thought to be rare (Nathan, 2006; but see Viana & al., 2015), but rarer still might be plant dispersals across long distances between different biomes (Crisp et al., 2009). The coincidence of intercontinental dispersal and biome shift, such as inferred in *Lupinus* (Drummond et al., 2012), *Bartsia* (Uribe-Convers & Tank, 2015), and *Hypericum* (Nürk, Michling & Linder, 2018), is assumed to be the result of a multifaceted interplay between geographical distance and ecological suitability of source and sink areas (Donoghue & Edwards, 2014). Here, we test the influence of these factors on the biogeographic history of the flowering plant genus *Erica* (Ericaceae).

The more than 800 *Erica* species across Europe and Africa provide an excellent example with which to test the impact of geographical and ecological distance on biogeographic history. Just 21 of the species are found in Central and Western Europe, Macaronesia, the Mediterranean and the Middle East. This species-poor assemblage nevertheless most likely represents the ancestral area of the clade (McGuire & Kron, 2005; Mugrabi de Kuppler et al., 2015; Kowalski & Fagúndez, 2017) where the oldest lineages began to diversify c. 30 Ma (Pirie et al., 2016). From around 15 Ma, a single lineage dispersed across different biomes of the Afrotemperate (sensu White, 1981): today 23 species are known from the high mountains of Tropical Africa; 51 in Southern Africa’s Drakensberg Mountains; c. 41 in Madagascar and the Mascarene islands; and c. 690 in the Cape Floristic Region of South Africa (Oliver, 2012; Pirie et al., 2016). Present day habitats of *Erica* species tend to be low nutrient and fire prone (Ojeda, 1998), but hence still differ markedly in ecology, from the Mediterranean climates of southern Europe and the Cape to colder climes of northern Europe and the non-seasonal temperate habitats of the high mountains in Tropical Africa. These habitats are also separated by considerable geographic distances, isolated by expanses of inhospitable ecosystems and/or ocean. Nonetheless, similar distribution patterns across Europe and Africa are observed in different plant groups (e.g. Galley et al., 2007; Gizaw et al., 2016).

Organisms adapted to different habitats respond differently to changing environmental conditions (Mairal, Sanmartín & Pellissier, 2017; Chala et al., 2017). For example, plant groups with greater tolerances of aridity than *Erica* may have had more contiguous past distributions across Africa (Bellstedt et al., 2012). Similar distribution patterns of such groups might thus be best described by biogeographic scenarios emphasising vicariance processes, such as for example the “Rand Flora”, representing plant lineages that show similar disjunct distributions around the continental margins of Africa (Sanmartín et al., 2010; Pokorny et al., 2015), or the “African arid corridor” hypothesis that seeks to explain disjunct distributions between the Horn of Africa and arid south-western Africa (Verdcourt, 1969; White, 1983). By contrast, similar distribution patterns observed across plants such as *Erica* that are adapted, or otherwise restricted, to habitats that remained largely isolated over time might instead be explained by concerted patterns of LDD (Knox & Palmer, 1998; Galbany-Casals et al., 2014; Nürk et al., 2015; Míguez et al., 2017). Examples include the shared arid adapted elements of Macronesia and adjacent North-West Africa and Mediterranean (Kim et al., 2008; Fernández-Palacios et al., 2011; García-Aloy et al., 2017), and the more mesic temperate or tropical alpine habitats of the “sky islands” of East Africa, in which, for example, multiple lineages originated from northern temperate environments (Gehrke & Linder, 2009; Gizaw et al., 2013, 2016).

A more specific biogeographic scenario, inferred from Cape clades with distributions very similar to that of *Erica,* involves dispersal north from the Cape to the East African mountains via the Drakensberg (“Cape to Cairo”; Galley & al., 2007). McGuire & Kron (2005) proposed a different scenario for *Erica* instead: southerly stepping stone dispersal through the African high mountains to the Cape. Both scenarios, however, imply that dispersal is more frequent between adjacent areas/over shorter distances. Short distance or stepping stone dispersal may indeed be more probable than LDD (Nathan, 2006), and distance alone could conceivably be more important than directionality (Linder et al., 2013). On the other hand, the probabilities of LDDs are hard to model (Higgins & Richardson, 1999; Nathan, 2006), in part because (observable) LDD events also involve successful establishment in more or less distinct environments (Donoghue & Edwards, 2014). Thus geographic distance and ecological suitability might individually constrain the biogeographic history of plants, or the interplay between both factors may be decisive (Donoghue, 2008; Carvajal-Endara et al., 2017), so much so that clades with similar ecological tolerances and origin might show convergence to similar distribution patterns (Merckx et al., 2015; Gizaw et al., 2016; Mairal, Sanmartín & Pellissier, 2017).

In this paper, we ask whether and to what extent geographic proximity or climatic niche similarity constrained the colonisation of the Afrotemperate by *Erica*. Until recent work (Pirie, Oliver & Bellstedt, 2011; Pirie et al., 2016), too little was known of the phylogenetic relationships of the 97% of *Erica* species outside Europe to be able to address such questions. Specifically, we test six biogeographic models, as illustrated in Fig. 1: Three that test the influence of geographic distance, climatic niche similarity, and the combination of both; and three differing stepping stone models that each imply geographical distance effects promoting dispersal predominantly between adjacent areas: northerly “Cape to Cairo”, “Southerly stepping stone” and a model that invokes elements of both, the “Drakenberg melting pot” hypothesis.

**Fig. 1:**
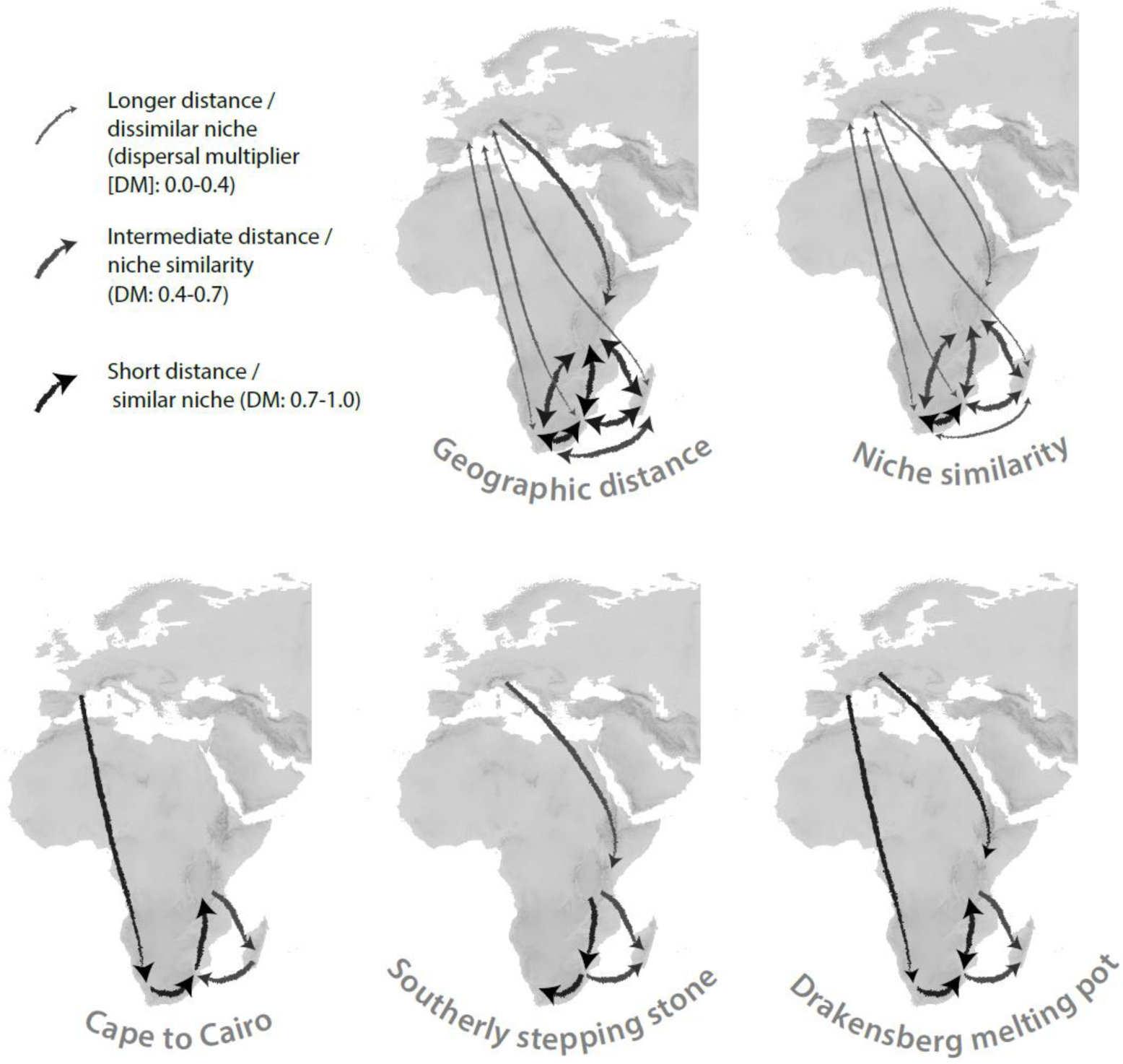
Biogeographic hypotheses. The pure geographic distance model; niche similarity, implying colonisation of areas with the most similar climatic niche (Donoghue 2008); these together constituting a combined geographic and niche similarity model; stepping stone: stepwise southerly colonisation of the Afrotemperate from Europe (following McGuire and Kron, 2005); Cape to Cairo: stepwise northerly colonisation of the Afrotemperate from the Cape (following Galley et al., 2007); necessarily preceded by LDD from Europe; and Drakensberg melting pot: the Drakensberg colonised by both southerly and northerly stepwise dispersal.

## Methods

### Phylogenetic hypothesis

Analyses were based on phylogenetic trees (Pirie et al., 2016; TreeBase study accession URL http://purl.org/phylo/treebase/phylows/study/TB2:S18291) which represent c. 60% of the c. 800 species of *Erica* from across their geographic range and DNA sequences from multiple plastid markers (*trnT-trnL* and *trnL-trnF-ndhJ* spacer sequences for all taxa, with exemplar sampling of *trnL* intron, *atpI-atpH* spacer, *trnK-matK* intron and *matK* gene, *psbM-trnH* spacer, *rbcL* gene, *rpl16* intron, *trnL-rpl32* spacer sequences) and nuclear ribosomal (nrDNA) internal transcribed spacer (ITS; for all taxa). For the biogeographic analyses here, we adopt the phylogenetic hypothesis of Pirie et al. (2016), the best tree inferred under Maximum Likelihood (ML) using RAxML (Stamatakis, 2006), based on the combined data and 597 taxa and rate smoothed using RELTIME (Tamura et al., 2012) with a single secondary calibration point derived from a wider fossil calibrated analysis of Ericaceae (Schwery et al., 2015). Pirie & al. (2016) identified a “Cape clade” that included all but one of the sampled species of *Erica* found in the CFR. The single exception was *E. pauciovulata*, which was placed within a polytomy including the Cape clade and other Afrotemperate lineages. This may, however, be artefactual due to sequence anomalies in the *trnL-trnF-ndhJ* spacer region of *E. pauciovulata*. Preliminary results based on additional sampling including nrDNA ETS (Pirie et al. in prep.) confirm the monophyly of Cape clade including *E. pauciovulata*, and we therefore exclude this taxon from biogeographic analyses to avoid inferring an independent colonisation of the CFR as a result of its uncertain position.

### Defining the pure-distance and the niche-based models

Five biogeographic areas of the *Erica* distribution were defined following Pirie & al. (2016): Europe (including northern Africa); Tropical Africa (TA); Madagascar; Drakensberg; Cape. We obtained occurrence data for *Erica* species from our own collections, and from PRECIS (representing mostly southern African collections, held by the South African National Biodiversity Institute; http://newposa.sanbi.org/) and GBIF (https://www.gbif.org/) databases. We curated the species occurrence data by removing obviously erroneous locality data, duplicated records, and records with imprecise occurrence data (coordinates with ≤ 3 decimal places, many of which represented centroids of quarter degree squares which were originally represented in PRECIS), but did not further consider the source of or information on the precision of the geographical coordinates, because these are most often not stated in the database-derived occurrence records. This resulted in 6818 individual occurrences representing the species in the phylogenetic trees (Appendix 1). Based on these individual occurrences we estimated the geographic range of the species in the five biogeographic areas (resulting in what we term ‘area ranges’, i.e. the distributional range of all of the species in a certain biogeographic area) We assessed the area ranges in a biogeographic area placing a buffer of one minutes of arc in radius (ca. 11 km) and 50 m elevation around the individual species occurrences (Europe 4667, Tropical Africa 42, Madagascar 70, Drakensberg 58, and Cape 1981 occurrences; Appendix 2) conservatively aiming at a representative approximation of spatial extent (Nakazato, Warren & Moyle, 2010; Anacker & Strauss, 2014) and ecological conditions of the species’ distribution in a respective biogeographic area. These area ranges, which include up to several thousands of spatial points, were used in the subsequent analyses to calculate geographical and ecological distances between biogeographic areas.

To incorporate a measure of geographic proximity among areas in a solely distance-based biogeographic model (the ‘geographic distance’ model; Fig. 1), we calculated the overall minimum geographic pairwise distances between the area ranges according to Meeus (1999) in WGS84 projection using the raster 2.3-33 package (Hijmans, 2015) in R (R Development Core Team, 2013). We converted geographic distances into dispersal rate multipliers (0-1, whereby the largest distance has the smallest dispersal probability), while comparing the effect of scaling the distances linearly (applying a linear model with intercept of 1 and a slope of −1.52^−07^) and exponentially (−0.25, −1 and −2).

To incorporate in a niche-based biogeographic model, the ‘niche similarity’ model (Fig. 1), a measure of climatic similarity between the biogeographic areas we built a multidimensional environmental model representing the full space of all available climates in the global study area (i.e. most of Europe and entire Africa, represented by >0.5 million spatially independently sampled point locations; Appendix 2) using principal component analysis (PCA) in R’s ade4 1.6-2 (Dray & Dufour, 2007). We then calculated the pairwise climate similarity between the area ranges (as defined by the species occurrence data; see above) as Schoener’s *D* (Schoener, 1968; ranging from 0 = no similarity to 1 = identical) while correcting for the entire climatic variance in the global study area and between the biogeographic areas (Broennimann et al., 2012) by kernel density smoothers using ecospat 2.1.1 (Broennimann et al., 2014). Finally, the biogeographic model for the niche similarity hypothesis was defined using the pairwise Schoener’s *D* values for the combined PCA axes 1 and 2 directly as dispersal rate multipliers between areas (for details see protocol in Appendix 2).

Finally, to consider both geographical and environmental distances in a joint model, also accounting for a negative correlation between both geographic and environmental distances (Kendall’s R = −0.64), we used two rate multiplier matrices, representing both climatic niche and physical distance (converted into probabilities; see above), as input.

### Biogeographic model testing and ancestral area reconstruction

We used BioGeoBEARS (Matzke, 2013) for parametric model testing, whilst aware of the debate surrounding these models and their comparison (Ree & Sanmartín, 2018; see Results and Discussion). The above defined biogeographic models (Fig. 1) were parameterized using different dispersal rate multipliers (see below and Appendix 3) and compared to null models that do not incorporate any constraints. As input data we used the rate-smoothed ML phylogeny reduced to one tip per sampled species (Pirie & al., 2016; the “best tree”), a file delimiting the distributional range of species, and a file indicating connectivity/distance between the different areas of the *Erica* distribution (varying for the different biogeographic models; Fig. 1, Appendix 3). Model fit of the different nested and non-nested models was tested using the Akaike Information Criterion (AIC) and the delta AIC (Burnham & Anderson, 2002). For model testing we additionally used nine trees from the RAxML bootstrap analyses of Pirie & al. (2016) of the same dataset (rate-smoothed using the ape package in R; Paradis & al., 2004). These trees were selected to represent the possible resolutions of phylogenetic uncertainty between the geographically restricted major clades (Appendix 4) but were otherwise chosen randomly with respect to topologies and branch lengths. All hypotheses were implemented with combinations of dispersal-extinction-cladogenesis (DEC; Ree et al., 2005; Ree & Smith, 2008), Bayarea-like or DIVA-like models, with or without allowing long distance dispersal (the “+J” model of Matzke, 2014). We focus on DEC and DEC+J models because these generally fit the data better than Bayarea-like or DIVA-like models.

Prior to comparing the different biogeographic hypotheses, we tested whether an unconstrained model fitted the data better than a) restricting the maximum number of areas at nodes to two; and/or b) implementing an adjacent area matrix (Appendix 3; Results). The Southerly stepping stone, Cape to Cairo, and Drakensberg melting pot hypotheses were then run under a range of different dispersal multipliers (0.00, 0.01, 0.05, 0.075, 0.1, 0.25 and 0.5; and for the DEC+J model also on the nine bootstrap trees with dispersal multipliers of 0.01, 0.1, 0.25 and 0.5) to test whether these arbitrary values influenced the results. For niche- and distance-based biogeographic models Schoener’s *D* values and geographic distances scaled to probabilities (see above) were used directly as dispersal multipliers (Appendix 3). We also assessed the impact on model fit (given the best fitting model) of a number of different values for the parameter “w”, which is an exponent for the dispersal multipliers (which otherwise was fixed to “1”; Appendix 3); and coding of *E. arborea* as European (following Mugrabi de Kuppler et al., 2015), rather than as widespread between Europe and Tropical Africa. Further details and example files for the BioGeoBEARS analyses are presented in Appendix 3.

### Estimating dispersal rates

For the best models under both DEC+J and DEC, given the best tree, we estimated the number and type of biogeographic events across the clade using Biogeographical Stochastic Modelling (BSM) as implemented in BioGeoBEARS (Matzke, 2014). BSM simulates histories of the times and locations of dispersal events. Frequencies were estimated by taking the mean and standard deviation of event counts from 50 BSMs. We also compared the results to that of simple parsimony optimisation using Mesquite v3.31 (Maddison & Maddison, 2006), under the assumption that LDD events are simply rare (Pirie et al., 2012). We incorporated phylogenetic uncertainty by summarising the results over the complete sample of 252 RAxML bootstrap trees adapted from Pirie & al. (2016), and coding *E. arborea* either as widespread between Europe and Tropical Africa or European (Appendix 5).

## Results

### Niche similarity model

The environmental space that represents all climates available in the study area – most of Europe and all of Africa – and that was used to approximate the climatic similarity between biogeographic areas (area ranges) calculated as hypervolume corrected Schoener’s *D* accounting for climatic variation in the study area and between the area ranges, explained >88% of the climate variation on the first two PCA axes. Despite the range of differing conditions within areas, e.g. with rainfall seasonality differing according to elevation, variation between area ranges was considerable (the distribution and the median values for Schoener’s *D* per PCA axis pairwise for the areas are presented in Appendix 6, and for the combined axes 1 and 2 in Appendix 7). According to the latter, the Cape and Drakensberg areas are climatically most similar (*D*: 0.71) and Europe and Madagascar are most different (*D*: 0.21). Most similar to the European are the Cape and Drakensberg climates (both *D*: 0.35), and the Tropical Africa climate (*D*: 0.27; Fig. 2C).

**Fig. 2:**
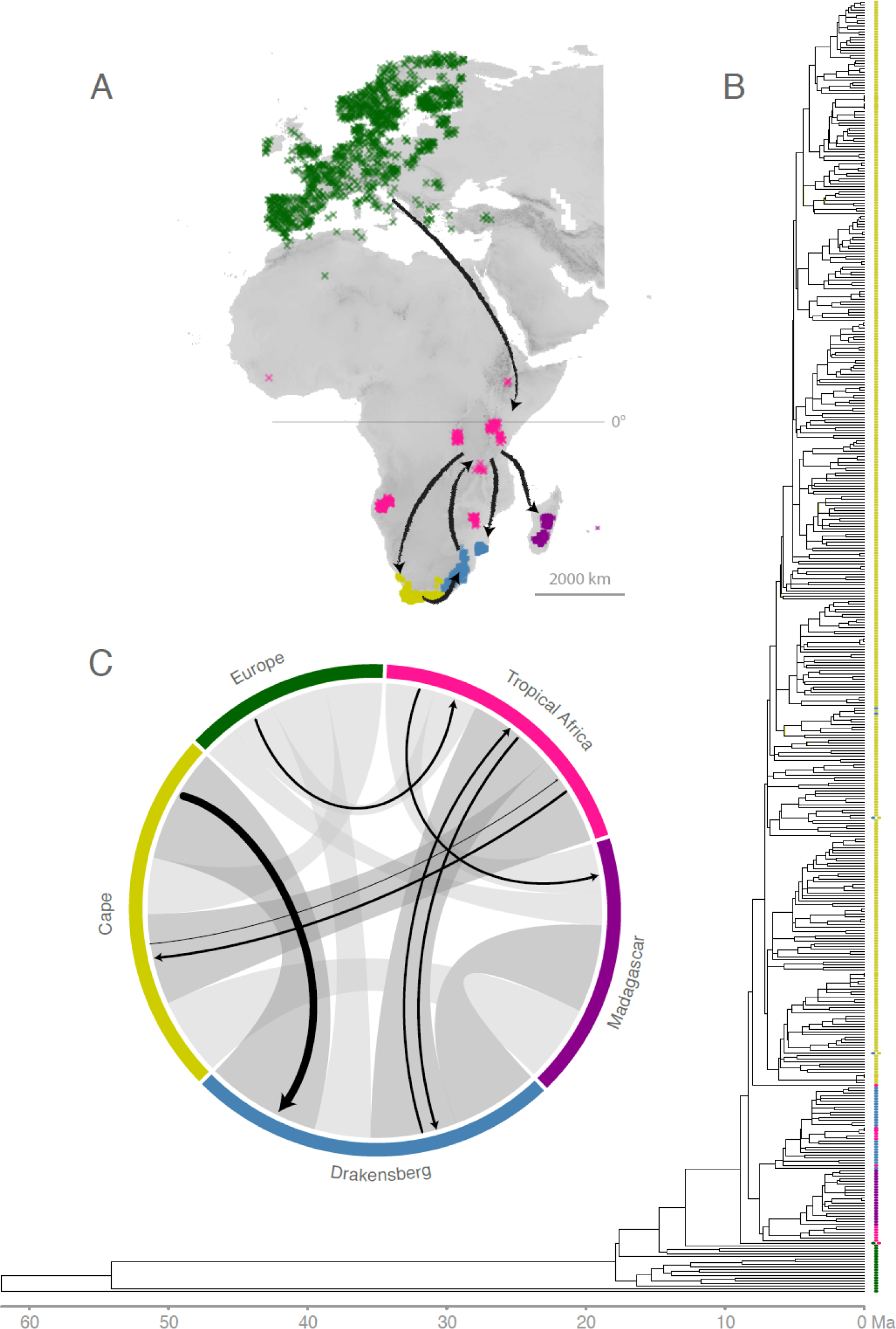
Biogeographic scenario. **A**) Inferred dispersal scenario between the biogeographic areas (colour coded “area ranges” as assessed by the buffered species occurrence data) depicted on the global study area. **B**) The phylogeny of *Erica* with a representation of ancestral areas derived from a single BSM analysis using the best tree and model (under DEC+J). **C**) Climate similarity between area ranges given as hypervolume corrected pairwise climate similarity (Schoener’s *D*; with *D* ≥ 0.5 in dark grey, and *D* < 0.5 in light grey; Table S8) with superimposed black arrows scaled to the dispersal rates per Ma inferred between areas (Table S13). Ma, million years.

### Biogeographic model testing

Assuming that AIC values of the differing models can be compared (but see Ree and Sanmartín, 2018), DEC/DEC+J models generally fit the data better than Bayarea-like or DIVA-like models and DEC+J models generally fit the data better than equivalent DEC ones (Appendix 8). Under DEC+J, models including an adjacent area matrix fitted the data better than those without constraint to dispersal. We additionally fixed the maximum number of ancestral areas to two, increasing the speed of the analyses without negatively impacting model fit. Under DEC, models with maximum areas at nodes restricted to two fitted the data better than those without constraint to ancestral ranges. Under both DEC+J and DEC, geographic distance fitted the data better when translated linearly into dispersal rate probabilities (0-1) than when scaled exponentially (Appendix 8); we therefore focus on models using the probabilities, referring to them simply as “geographical distance”. The DEC+J results in general do not show the flaws as reported by Ree and Sanmartín (2018). For example, the values for range expansion (parameter d) were similar and low (0.0030 and 0.0027 per Ma respectively; Appendix 9). Under DEC+J, cladogenetic dispersal (parameter j) was 0.0024 per node, i.e. lower than d (particularly given an average branch length across the *Erica* phylogeny of 1.78 Ma, variance of 11.67) and much lower than the maximum permitted value (3).

Under DEC+J given the best tree, the “Drakensberg melting pot”, “geographic distance”, and “Southerly stepping stone” models revealed the best fit (lowest AIC with deltaAIC ≥2); under DEC the Drakensberg melting pot model alone scored best, but with AIC 141 compared to AIC 131 for DEC+J (Appendix 8). Adopting DEC+J as the generally better fitting and biologically more realistic model (see Discussion), we assessed the results given phylogenetic uncertainty represented by selected bootstrap trees. Based on the bootstrap trees, the combined niche-geographic distance hypothesis was often among the best fitting models (deltaAIC <2 given eight of nine trees), scoring better than pure distance (deltaAIC <2 for five trees), or niche similarity (deltaAIC <2 for four trees) alone. The “Cape to Cairo” model generally fitted better than most other biogeographic scenarios (deltaAIC <2 for eight of nine trees, compared to Drakensberg melting pot (deltaAIC <2 for two of nine trees) and southerly stepping stone (not amongst the best fitting models); Table 1; Appendix 8).

**Table 1:** Best fitting biogeographic models given the best tree (DEC+J and DEC) and nine selected bootstrap trees (DEC+J). Dispersal multipliers are indicated where relevant, as are the Log likelihood (LnL), Akaike Information Criterion (AIC), and overall deltaAIC scores for models. Models with deltaAIC of 0 are indicated in bold type. DMP=Drakensberg mekting-pot; Dist=Distance; SSS=Southerly stepping-stone; CtoC=Cape to Cairo.

| Tree | Model | Dispersal multiplier | LnL | AIC | deltaAIC |
| --- | --- | --- | --- | --- | --- |
| **Best (DEC+J)** | **DMP** | **0.5** | **-62.5** | **131** | **0** |
|  | **DMP** | **0.1** | **-62.5** | **131** | **0** |
|  | **DMP** | **0.25** | **-62.5** | **131** | **0** |
|  | DMP | 0.75 | -62.6 | 131.2 | 0.2 |
|  | Dist | - | -62.8 | 131.6 | 0.58 |
|  | SSS | 0.5 | -63.1 | 132.3 | 1.3 |
|  | SSS | 0.25 | -63.3 | 132.7 | 1.7 |
| **Best (DEC)** | **DMP** | **0.75** | **-68.6** | **141.2** | **0** |
|  | DMP | 0.5 | -68.7 | 141.3 | 0.1 |
| **BS 0_0** | **Niche+Dist** | **-** | **-61.2** | **128.4** | **0** |
|  | CtoC | 0.25 | -61.7 | 129.4 | 1 |
|  | CtoC | 0.1 | -62.1 | 130.1 | 1.7 |
|  | Dist | - | -62.2 | 130.4 | 2 |
| **0_1** | **Niche+Dist** | - | **-65.6** | **137.1** | **0** |
|  | CtoC | 0.1 | -65.8 | 137.6 | 0.5 |
|  | CtoC | 0.25 | -66.1 | 138.3 | 1.2 |
| **0_2** | **Niche+Dist** | - | **-60.2** | **126.3** | **0** |
|  | CtoC | 0.25 | -60.4 | 126.7 | 0.4 |
|  | CtoC | 0.1 | -60.4 | 126.9 | 0.6 |
|  | Dist | - | -61 | 127.9 | 1.6 |
| **1_0** | **CtoC** | **0.1** | **-58.7** | **123.5** | **0** |
|  | CtoC | 0.25 | -59.5 | 124.9 | 1.4 |
| **1_1** | **Niche+Dist** | - | **-61.7** | **129.4** | **0** |
|  | Niche | - | -62.6 | 131.2 | 1.8 |
| **1_2** | **Niche+Dist** | - | **-55.9** | **117.9** | **0** |
|  | CtoC | 0.25 | -56.5 | 119 | 1.1 |
|  | CtoC | 0.1 | -56.9 | 119.7 | 1.8 |
|  | Dist | - | -56.9 | 119.7 | 1.8 |
| **2_0** | **Niche+Dist** | - | **-62.3** | **130.6** | **0** |
|  | CtoC | 0.25 | -62.6 | 131.2 | 0.6 |
|  | Dist | - | -62.8 | 131.5 | 0.9 |
|  | CtoC | 0.1 | -62.9 | 131.8 | 1.2 |
|  | CtoC | 0.5 | -63.1 | 132.3 | 1.7 |
|  | Niche | - | -63.2 | 132.4 | 1.8 |
| **2_1** | **Dist** | - | **-56.5** | **119.1** | **0** |
|  | DMP | 0.1 | -56.6 | 119.2 | 0.1 |
|  | DMP | 0.25 | -56.6 | 119.2 | 0.1 |
|  | DMP | 0.5 | -57 | 120 | 0.9 |
|  | CtoC | 0.5 | -57.2 | 120.4 | 1.3 |
|  | Niche+Dist | - | -57.2 | 120.4 | 1.3 |
|  | CtoC | 0.25 | -57.5 | 121 | 1.9 |
|  | Niche | - | -57.6 | 121.1 | 2 |
| **2_2** | **Dist** | - | **-65.3** | **135.6** | **0** |
|  | Niche+Dist | - | -65.2 | 136.4 | 0.8 |
|  | DMP | 0.25 | -65.5 | 137 | 1.4 |
|  | CtoC | 0.5 | -65.5 | 137 | 1.4 |
|  | DMP | 0.1 | -65.5 | 137.1 | 1.5 |
|  | CtoC | 0.25 | -65.6 | 137.1 | 1.5 |
|  | Niche | - | -65.8 | 137.5 | 1.9 |
|  | DMP | 0.5 | -65.8 | 137.6 | 2 |

### Ancestral area reconstruction

Overall, we infer a colonisation path of *Erica* from Europe to the Cape via an initial migration to Tropical Africa, under DEC+J and irrespective of best fitting model or phylogenetic uncertainty. When *E. arborea* is treated as widespread between Europe and Tropical Africa, the common ancestor of the African/Madagascan clade is inferred to have been similarly widespread. When *E. arborea* is treated as ancestrally European, dispersal from Europe to Tropical Africa is inferred without a transitional widespread distribution. Under DEC, the colonisation path to the Cape is also via an initial migration to Tropical Africa, then a widespread distribution between Tropical Africa and the Cape, followed by an extinction in Tropical Africa. Whether *E. arborea* is treated as widespread between Europe and Tropical Africa or not, the common ancestor of the African/Madagascan clade is inferred to have been similarly widespread between Europe and tropical Africa. Ancestral area reconstructions given the best tree under the best fitting models (as well as under a model without range or dispersal constraints for comparison; in each case under both DEC+J and DEC) are presented in Appendix 13. Overall, ancestral areas inferred under parsimony were consistent with those inferred under parametric models (more so with those under DEC+J, given that widespread distributions are not incorporated into standard character optimisation), with the numbers and directions of shifts unaffected by phylogenetic uncertainty.

The vast majority of biogeographic events inferred using BSM under both DEC+J and DEC were within-area speciation (97.15 % and 96.26% respectively; Appendix 9). Under DEC+J, few range expansion events were inferred between Europe and Tropical Africa and between Tropical Africa and the Drakensberg region, with most between Cape and the Drakensberg regions (Appendix 10). Dispersal rates between area ranges inferred under BSM are summarised in Fig. 2C. A single founder event (parameter j) was inferred from Tropical Africa to the Cape region, with fewer events between the Drakensberg and Tropical Africa and between Tropical Africa and Madagascar. Overall, most founder events took place from Tropical Africa (1.96 [standard deviation of 0.47] events averaged across 50 BSM; Appendix 11). In addition to the most commonly inferred range expansions given DEC+J, under DEC additional range expansions were inferred from Tropical Africa to Madagascar and from Tropical Africa to the Cape (Appendix 10). With each range expansion under DEC, the corresponding ancestral distribution was widespread. Under both DEC+J and DEC dispersal rates between Tropical Africa and the Drakensberg were roughly symmetrical, as opposed to those between the Cape and the Drakensberg or between Europe and Tropical Africa which were asymmetrical (Fig. 2; Appendix 12).

## Discussion

In this study, we modelled shifts between biomes and dispersals over larger distances in the evolution of *Erica*, in order to test six hypotheses for the origins of Afrotemperate plant groups (Fig. 1). Three models concerned general factors considered of importance in limiting plant dispersal: geographical distance, similarity of realised climatic niches, and a combination of geographical and ecological proximity. The remaining three models described specific colonisation hypotheses of the Afrotemperate, in each case proposing a stepwise shift in distributions between adjacent areas. These models differed in the area of origin and in the direction of dispersal: northerly dispersal from the Cape (“Cape to Cairo”), versus southerly dispersal from Europe (“Southerly stepping stone”), or a combination of both (termed here “Drakensberg melting-pot”).

Of the stepping-stone-dispersal models, “Cape to Cairo” and/or “Drakensberg melting-pot” fit the data better than “Southerly stepping stone” for all but the best tree, but relative fit of the models was somewhat sensitive to phylogenetic uncertainty (Table 1). By contrast, the positions of areas relative to one another, and the similarities in their realised climatic niche, were consistently prominent in our results. Of the distance models, the combination of geographical and ecological distance fit the data well. Our results showed that these factors are correlated across the *Erica* distribution, but nevertheless given the phylogenetic uncertainty it was the combination of both that often fitted the data better than either of factor individually (or indeed the stepping stone models). The generally better fit of the combined geographic and realised niche model affirms the concerted importance of both factors in shaping distributional patterns of plants (Donoghue 2008; Donoghue and Edwards, 2014). Of the nine range expansion events that we inferred (DEC+J, best tree, best model), seven respectively were between adjacent areas or between areas with similar environmental conditions (where “similar” is arbitrarily defined as a pairwise Schoener’s *D* > 0.5; Fig. 2). Overall, this represents striking evidence for geographical and ecological distance constraining past and present distributions of *Erica* species, similar to that inferred for other Mediterranean climate plant groups (Skeels & Cardillo, 2017). Irrespective of model fit, the sequence of dispersal events that we inferred from ancestral area reconstructions, based on both the set of best fitting parametric models and a parsimonious interpretation of the infrequent dispersal events (Fig. 2), does resemble a “Drakensberg melting-pot” scenario. The Drakensberg acted as a sink for dispersals from the adjacent Cape and Tropical African regions, but not as a stepping stone (or indeed a “springboard”; Galley & al., 2007).

Cape lineages found in the Drakensberg have not dispersed to Tropical Africa, and neither have Tropical Africa lineages found in the Drakensberg dispersed further to the Cape. This is unexpected, not only because of the low distances and high niche similarities involved, but also because of the equivalent events inferred in other similarly distributed plant groups (Galley et al., 2007). Striking in a different way are three unique events: the single dispersals from Europe to Tropical Africa, out of Tropical Africa to the Cape, and out of Tropical Africa to Madagascar, which were each over much longer distances. The dispersals to Tropical Africa and to Madagascar both might have involved shifts in realised niches (indicated by low Schoener’s D values of 0.298 and 0.274 respectively); that to the Cape, borderline so (Schoener’s D of 0.560; Fig. 2). Notably, the dispersals to tropical Africa and to the Cape coincided with clear increases in diversification rate (Pirie et al., 2016).

Potential explanations for these apparent exceptions to the general importance of geographical and ecological distance might be found in the context of the changing climates and geology of the African continent during the timeframe of the *Erica* radiation. The summer-arid climate of the present day Cape has been linked to the establishment of the cold Benguela current off the south-west African coast in the mid Miocene 14-10 Ma (Marlow et al., 2000; Zachos et al., 2001). Evidence from pollen deposited in nearby marine sediments shows an accumulation of typical Cape lineages since roughly the same time, including Ericaceae (Dupont et al., 2011), supported by further evidence from recent dated phylogenies both for the ages of clades in the Cape (e.g. Verboom & al., 2009; Hoffmann & al., 2015) and the origins of fire adapted lineages (Bytebier et al., 2011). The gradual change from a more tropical to a mesic flora and initiation of a regular fire regime in south-western Africa might be ecological changes important for the establishment of *Erica* in the Cape. Whilst the mountains of the Western Cape, home to much of the *Erica*-dominated fynbos vegetation, long predate Miocene climatic changes, the origins of the Drakensberg and Tropical African high mountains, *Erica’s* area of first establishments in Africa, are more recent, with uplift in these regions creating montane habitats from the Miocene onwards (McCarthy & Rubidge, 2005).

Thus, shifting climates and mountain building created an archipelago of temperate islands across sub-Saharan Africa that were available for colonisation by plants able to tolerate the novel conditions. These included *Erica* species, which had begun to diversify c. 30 Ma in the Northern Hemisphere (Pirie et al., 2016), and which as a clade are characterised by drought resistant leaves (Van Wilgen, Higgins & Bellstedt, 1990) as well as adaptations to post-fire regeneration (Ojeda, 1998). However, our analyses of the realised climatic niches of *Erica* species in their different biomes suggests that despite these pre-existing drought and fire adaptations, colonisation of new areas by *Erica* involved further adaptation to rather different climatic conditions, as inferred for tropical alpine *Hypericum* in South America (Nürk, Michling & Linder, 2018) and hypothesised for tropical high alpine plants in general (Gehrke, 2018). In this context, biome shifts and increased diversification rates may be linked: the open field presented by these newly formed, isolated, temperate habitats may have facilitated both the chance establishment of suboptimally adapted plants and their subsequent *in situ* shift into new adaptive zones, promoting accelerated diversification.

Neither differences in ecological nor geographic distance present an obvious explanation for why dispersal to the Drakensberg was not followed by further independent colonisations, particularly of the Cape. One possibility could be that within the Drakensberg, Cape and Tropical African elements occupy somewhat differing niches: the former, such as the widespread Cape-Drakensberg species *E. cerinthoides* and *E. caffra* predominantly at lower elevations, the latter at higher elevations under conditions differing more to those in the Cape. Another could be niche pre-emption (Silvertown, 2004), whereby the single colonisation and rapid species radiation of Cape *Erica* prevented further colonisation by similar competitors.

Widespread species such as *E. cerinthoides* and *E. caffra*, found in the Cape and Drakensberg, and *E. arborea*, found in Europe and Tropical Africa (Désamoré & al., 2011; Gizaw & al., 2013), are exceptional in *Erica*. Almost all extant species are restricted to just one of the areas as defined here (including species such as *E. silvatica* and *E. benguelensis* that are widespread across disjunct areas within Tropical Africa), and species radiations leading to most of the present day diversity of *Erica* were within single areas (Pirie & al., 2016). This suggests that species distributions were restricted throughout the evolution of the *Erica* African/Madagascan clade, and that the areas remained isolated during this period (i.e. the last c. 15 Ma; Pirie & al., 2016). We would also argue that it lends credibility to results obtained under DEC+J, in which some range shifts were treated as cladogenetic dispersal events (instead of by inferring seemingly implausible widespread distributions), despite arguable drawbacks in the implementation of that model (Ree & Sanmartín, 2018).

Nevertheless, the extent and position of suitable habitats across the Afrotemperate shifted considerably during this timeframe, and the implicit assumption of our analyses, that they can be treated as consistent during the *Erica* diversification, is a considerable simplification. This may not impact the results of a broader scale analysis such as the one we present here, but might be important in the context of diversifications within regions, such as those within the Cape (Linder, 2003; Dupont et al., 2011; Schnitzler et al., 2012; Hoffmann, Verboom & Cotterill, 2015) or Drakensberg (Bentley, Verboom & Bergh, 2014). To assess the impact of climatic changes on the dramatic radiation of Cape *Erica*, for example, it would be important to translate realised niches into past distributions to model the shifting extents and interconnectedness of populations through time (cf. Mairal & al., 2017).

### Conclusion

The overall picture to be gleaned from the colonisation history of *Erica* across the Afrotemperate is one of infrequent dispersal limited by geographic distance and ecological similarity. Lack of dispersals where they might be expected – in the case of *Erica*, the Drakensberg acting as a sink, rather than stepping stone to wider dispersal – can point to biological and historical idiosyncrasies of particular lineages. Our results also show the importance of single unique events that can run counter to general trends. In *Erica*, three particularly long distance dispersals, two potentially with shifts in the realised niche, were followed by species radiations – most notably in the Cape – that dominate the narrative of the group as a whole. Our results serve to further emphasise the importance of such rare events, in which unique biome shifts fuel dramatic evolutionary radiations.

## Acknowledgements

This preprint has been reviewed and recommended by Peer Community In Evolutionary Biology (https://dx.doi.org/10.24072/pci.evolbiol.100065). Invaluable constructive comments on previous versions of the preprint were provided by Andrea Meseguer, Simon Joly, Florian Boucher, and two anonymous reviewers. We thank J. Fagúndez, A. Hitchcock, R. Turner, M. Muasya, C. Stirton, R. Clark, B. Bytebier, M. Pimentel, F. Ojeda, C. Merry, and many others for providing samples and Cape Nature and South Africa National Parks for assistance with permits. We also gratefully acknowledge the computing time granted on the supercomputer Mogon at Johannes Gutenberg University Mainz (www.hpc.uni-mainz.de), and F. Michling for providing R code. Funding was provided by the South African National Research Foundation (NRF; DUB and MDP); a postdoctoral fellowship from the Claude Leon Foundation (MDP); DFG (PI1169/1-1, PI1169/1-2, PI1169/2-1 and PI1169/3-1 to MDP); and the Ministerium für Klimaschutz, Umwelt, Landwirtschaft, Natur-und Verbraucherschutz des Landes Nordrhein-Westfalen, the Faculty of Agriculture Lehr-und Forschungsschwerpunkt “Umweltverträgliche und Standortgerechte Landwirtschaft”, Bonn University; and the Landgard foundation (AMK). Any opinion, finding and conclusion or recommendation expressed in this material is that of the authors and the NRF does not accept liability in this regard.

## Appendices

Appendix 1: Methods: occurrence data

Appendix 2: Methods: Global environmental space, area ranges, and climate similarity analysis

Appendix 3: Methods: Biogeographic models; example files for BioGeoBEARS analyses

Appendix 4: Methods: Selected bootstrap trees used to represent phylogenetic uncertainty between geographically restricted major clades

Appendix 5: Methods: Mesquite file used for parsimony ancestral state reconstruction including RAXML bootstrap trees

Appendix 6: Results: pairwise climate similarity (Schoener’s *D*) between biogeographic areas per PC axis

Appendix 7: Results: Pairwise climate similarity (Schoener’s *D*) between biogeographic areas for combined PC axes

Appendix 8: Results of the different models under DEC+J and DEC (generally the better models compared to DIVA-like and BAYAREA-like-models)

Appendix 9: Results: Summary of event counts from 50 biogeographical stochastic mappings under the best inferred model using the best tree

Appendix 10: Results: Number of range-expansion dispersal events (mean and standard deviation of all observed “d” dispersals) averaged across 50 biogeographical stochastic mappings under the best inferred model using the best tree.

Appendix 11: Results: Number of cladogenetic dispersal events (mean and standard deviation of all observed jump ‘j’ dispersals) averaged from 50 biogeographical stochastic mappings under the best inferred model using the best tree.

Appendix 12: Results: Number of all dispersal events (mean and standard deviation of all observed anagenetic ‘a’, ‘d’ dispersals, PLUS cladogenetic founder/jump dispersal) averaged from 50 biogeographical stochastic mappings under the best inferred model using the best tree.

Appendix 13: Results: Ancestral area reconstructions inferred using BioGeoBEARS given the best tree under the best fitting model given A: DEC+J; B: DEC; and without range or dispersal constraint: C: DEC+J; D: DEC. For each model the single most probable state is shown first (boxes with areas at nodes) followed by the relative probability of each state represented with pie charts at nodes. Areas are represented by colours: Dark blue for Europe (E); green for Tropical Africa (T); yellow for Madagascar (M); light blue for Drakensberg (D); red for Cape (C); and further colours for widespread distributions as indicated in the legends.

