## Appendix 2 for "Leaps and bounds: geographical and ecological distance constrained the colonisation of the Afrotemperate by *Erica*"

**Appendix 2: Methods: Global environmental space, area ranges, and climate similarity analysis**

1) To construct a global environmental space we sampled ≥0.5 million spatially independent point locations in Africa and Europe defining the “global study area” (i.e. the range within which all species in this study are found) by a polygon with ymax = 74, ymin = -40, xmax = 60, xmin = -32 adjusted to geographic distribution of the *Erica* species occurrences used in this study (Fig. S2.1). Based on species occurrences (Appendix 1) we defined the spatial and altitudinal max/min range of the biogeographic areas obtaining a spatial sample of ‘background ranges’ (Table S2.1; Fig S2.1) used to correct for regional differences in the available climates (Broennimann & al., 2012).

**Table S2.1.** Definition of ranges detailing geographic and elevational extent.

| **Background Range** | **y-max** | **y-min** | **x-max** | **x-min** | **alt-max** | **alt-min** |
| --- | --- | --- | --- | --- | --- | --- |
| Europe (E) | 72.01 | 27.10 | 42.82 | -29.65 | 2580 | -55 |
| Tropical Africa (T) | 8.67 | -20.79 | 40.82 | -9.37 | 4225 | 1091 |
| Madagascar (M) | -17.91 | -23.64 | 58.47 | 44.34 | 2561 | 551 |
| Drakensberg (D) | -23.49 | -33.56 | 34.97 | 25.96 | 2835 | -1 |
| Cape (C) | -28.99 | -35.83 | 27.49 | 16.44 | 2196 | -50 |

‘Area ranges’ were defined around species occurrences by sampling per biogeographic area all point locations within a buffer of ±1 minutes of arc lat/long and ±50 m elevation around single species occurrences aiming at a conservative approximation of species ranges and the associated climatic conditions (Figs. S2.1, S2.2). These area ranges were also used to calculate the minimum distances used to define the dispersal multipliers (the dispersal probabilities between areas) in the ‘pure-distance’ biogeographic model. Coordinates of area ranges are presented in Appendix 1 (file “Appendix1_area_data.txt”).

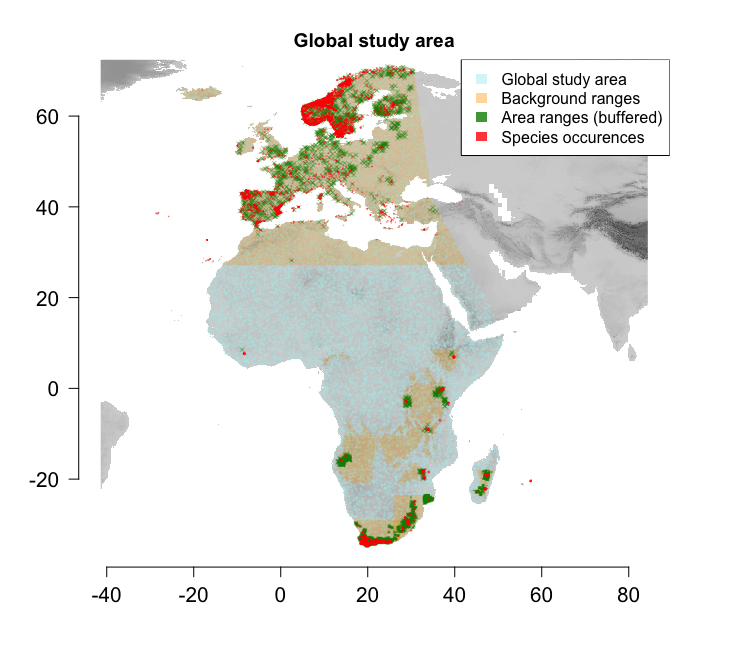

**Fig. S2.1.** Global study area, background ranges, area ranges, and species occurrences used to approximate climatic similarity between biogeographic areas. Background ranges per biogeographic area are defined by min/max species occurrences per area. Area ranges represent buffered ranges around single species occurrences.

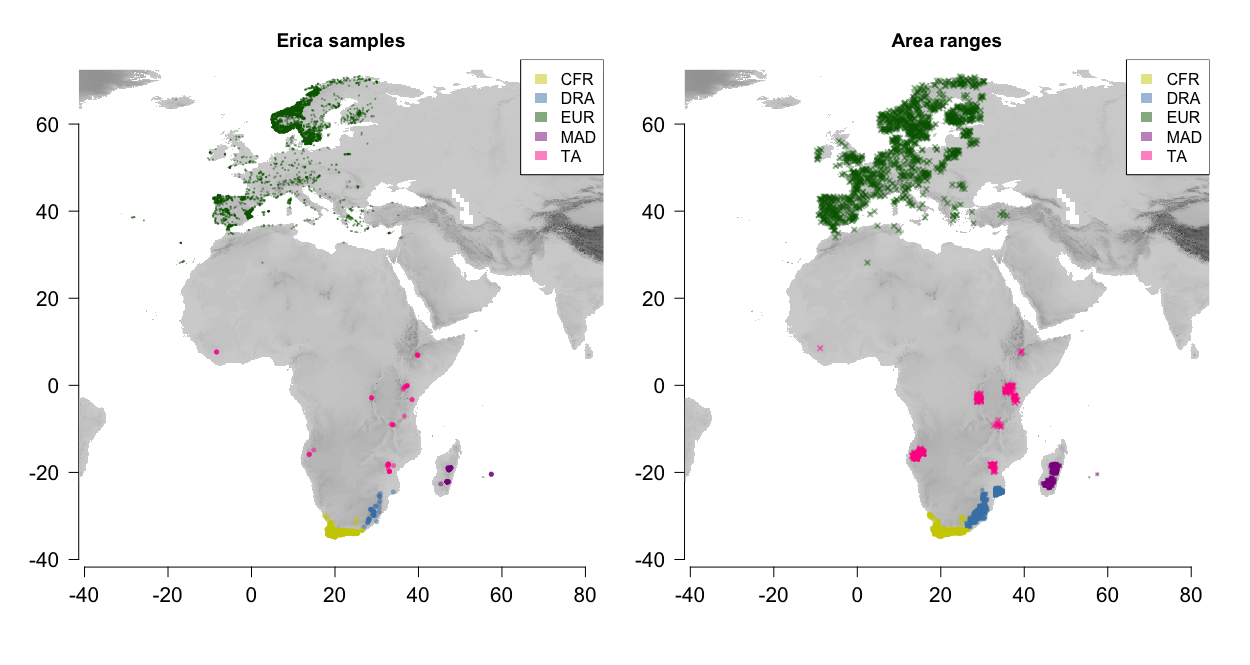

**Fig. S2.2.** Species occurrences (to the left) and area ranges (buffered around species occurrences; to the right) coloured according to biogeographic area (legend top right). CFR, Cape; DRA, Drakensberg; EUR, Europe; MAD, Madagascar; TA, Tropical Africa.

2) We then extracted climate data from the WorldClim version 1.3 BioClim data with a spatial resolution of 30’ (Hijmans et al., 2005) corresponding to spatial points and species occurrences using raster 2.3-33 in R (R Development Core Team, 2013; Hijmans, 2015). We accounted for collinearity in the BioClim data by removing variables that had high pairwise correlation to most others (Pearson’s *R* ≥ 7.5, calculated in R’s default stats package) retaining six variables (Table S2.1), which were used to construct the environmental space. We tested for spatial autocorrelation among the six selected environmental variables within the global study area as a function of distance using a Mantel correlogram calculated in R using ecospat 2.1.1 (Broennimann et al., 2014). We defined 1000 km as the maximum distance to be computed in the correlogram, using 10 classes of distances, and 100 permutations to calculate significance. No significant spatial autocorrelation was detected in the data (Fig. S2.3). Climate data of species occurrences and area ranges are presented in Appendix [1] in the extra SI files “Appendix1_Erica_data.txt” and “Appendix1_area_data.txt”.

**Table S2.2.** Climate variables used to construct the environmental space. Contribution of variables to PCA axes are detailed by column coordinates (BioClim, number of BioClim variable).

| **Variable** | | **BioClim** | **Axis 1 (56.37%)** | **Axis 2 (31.79%)** |
| --- | --- | --- | --- | --- |
| **Temperature** [°C] | |  |  |  |
|  | Mean annual temperature | 1 | -0.9338695 | 0.2757208 |
|  | Mean diurnal range | 2 | -0.8669087 | -0.2425715 |
|  | Temperature seasonality | 4 | 0.2955829 | -0.9047143 |
|  | Mean temperature of coldest quarter | 11 | -0.871166 | 0.4386694 |
| **Precipitation** [mm] | |  |  |  |
|  | Annual precipitation | 12 | 0.4759796 | 0.7787043 |
|  | Precipitation of driest quarter | 17 | 0.8281957 | 0.3937771 |

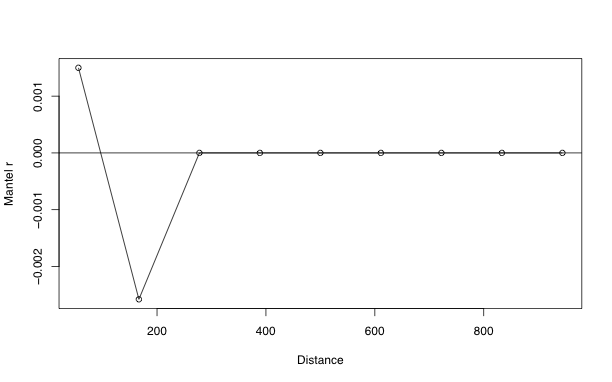

**Figure S2.3**. Mantel correlogram on environmental variables. The Mantel r represents the dissimilarity in variable composition and distance is in km. Note that the graph indicates no spatial autocorrelation significantly different from zero among the six selected bioclim variables used to construct the environmental space.

3) We finally constructed the environmental space using principal component analysis (PCA) in the R package ade4 1.6-2 (Dray & Dufour, 2007) after log-transforming the variables to produce equal spreads. The first two principal components (PC axes) were selected for the subsequent analysis based on the broken-stick criterion (Jackson, 1993).

4) To obtain a measure of climate similarity between the biogeographic areas we calculated pairwise hypervolume corrected Schoener’s *D* per PC axis in R using ecospat 2.1.1 (Broennimann et al., 2014) a thousand times, each time sampling a thousand points per area range. The final pairwise similarity between areas was calculated as the combined median of PC axes 1 and 2, which were directly used as dispersal multipliers (i.e. defining the dispersal probabilities between the areas) in the niche-based biogeographic model.

R Development Core Team. 2013. R: A language and environment for statistical computing.
