## Appendix 3 for "Leaps and bounds: geographical and ecological distance constrained the colonisation of the Afrotemperate by *Erica*"

**Appendix 3: Methods: Biogeographic models; example files for BioGeoBEARS analyses**

Prior to comparing the different biogeographic hypotheses we tested whether an unconstrained model fitted the data better than a) restricting the maximum number of areas at nodes to two; and/or b) implementing an adjacent area matrix. Both a) and b) would be consistent with the present day distributions of the species, which never exceed two, adjacent, areas. Even distributions widespread across just two adjacent areas are rare, and given the geographic structure in the *Erica* phylogeny we would assume that this was the case throughout the time of the radiation (Pirie et al., 2016). The adjacent area matrix in particular explicitly disallows reconstructions of widespread yet interrupted distributions at ancestral nodes. We would argue that such distributions are unlikely in principle and that dispersal between such areas – if infrequent – would be more appropriately modelled by a process equivalent to jump dispersal than by assuming ongoing gene flow. For DEC+J an adjacent area matrix fit the data better (Results), and we therefore implemented it in subsequent models. Restricting maximum areas to two, by contrast, had a negligible impact on model fit (Results); to increase the speed of the analyses we nevertheless restricted maximum (ancestral) areas to two in subsequent models, thereby assuming that in the past *Erica* species also never occurred across more than two regions. For DEC, by contrast, the adjacent area matrix did not improve model fit (and was therefore not further implemented), but restricting ancestral areas to two did (and was). The best fitting model was then used as basis and compared to models implementing the different dispersal hypothesis.

The parameter “w”, which is an exponent for the dispersal multipliers, was fixed to “1”. A free “w” parameter can help to remove some of the subjectivity of dispersal multipliers by optimising the matrix but there are known issues in finding the optimal maximum likelihood using a free “w” parameter ([phylo.wikidot.com/biogeobears#toc20](http://phylo.wikidot.com/biogeobears#toc20)). We also tried different values for “w” using only the best tree and the most likely dispersal model (Drakensberg melting pot with a multiplier of 0.25 for DEC+J and with a multiplier of 0.075 for DEC) to test the effect. The differences in model fit were negligible (DEC+J) or negative (DEC respectively). We therefore only report results fixing “w” to 1.

Mugrabi de Kuppler et al. (2015) inferred reticulation in Europe between ancestors of *E. lusitanica* and *E. arborea* on the basis of gene tree incongruence. This incongruence led Pirie & al. (2016) to exclude *E. lusitanica* from the combined analyses of plastid and nuclear DNA sequences used here, whereby the evidence that it presents for a European origin of *E. arborea* was inevitably lost from these analyses. We therefore ran the best model coding *E. arborea* as European, rather than as widespread between Europe and TA, to test the potential impact on our results of a European ancestral area for the species*.*

We also tested models that solely incorporates distances (either physical or environmental), based on the best model using the initial constraints (adjacency matrix and/or maximum area set to two: DEC+J/DEC respectively). For the pure distance model, we tested distances converted to probabilities, a linear model and exponentially transformed distances (0 to1). To combine both geographical and environmental distances in a joint model, instead of a single rate multiplier matrix both niche-based and distance-based matrices were used.

**Table S3.1**. Biogeographic models, here shown with 0.00 dispersal multipliers for unlikely dispersal pathways. Abbreviations: E, Europe; D, Drakensberg; T, Tropical Africa; M, Madagascar; C, Cape. Note that for the Stepping Stone, Cape to Cairo and Drakensberg Melting Pot model we used different dispersal multipliers: 0.00, 0.01, 0.05, 0.075, 0.1, 0.25 and 0.5. Here we only show the 0 matrices.

| Model | Areas | E | T | M | D | C |
| --- | --- | --- | --- | --- | --- | --- |
| Stepping Stone | E | 1 | 1 | 0 | 0 | 0 |
|  | T | 0 | 1 | 1 | 1 | 0 |
|  | M | 0 | 0 | 1 | 0 | 0 |
|  | D | 0 | 0 | 1 | 1 | 1 |
|  | C | 0 | 0 | 0 | 0 | 1 |
| Cape to Cairo | E | 1 | 0 | 0 | 0 | 1 |
|  | T | 1 | 1 | 1 | 0 | 0 |
|  | M | 0 | 0 | 1 | 0 | 0 |
|  | D | 0 | 1 | 1 | 1 | 0 |
|  | C | 0 | 0 | 0 | 1 | 1 |
| Drakensberg melting pot | E | 1 | 1 | 0 | 0 | 1 |
|  | T | 1 | 1 | 1 | 1 | 0 |
|  | M | 0 | 0 | 1 | 0 | 0 |
|  | D | 0 | 1 | 1 | 1 | 0 |
|  | C | 0 | 0 | 0 | 1 | 1 |
| Geographic distance |  |  |  |  |  |  |
| in m | E | 0 | 2480098 | 6411884 | 6482646 | 6570434 |
| (reference | T | 2480098 | 0 | 1208094 | 404243 | 1380045 |
| only) | M | 6411884 | 1208094 | 0 | 984007 | 2059502 |
|  | D | 6482646 | 404243 | 984007 | 0 | 52129 |
|  | C | 6570434 | 1380045 | 2059502 | 52129 | 0 |
| as “Probability” | E | 1 | 0.623 | 0.025 | 0.014 | 0.001 |
| (0-1) | T | 0.623 | 1 | 0.816 | 0.939 | 0.790 |
|  | M | 0.025 | 0.816 | 1 | 0.850 | 0.687 |
|  | D | 0.014 | 0.939 | 0.85 | 1 | 0.992 |
|  | C | 0.001 | 0.790 | 0.687 | 0.992 | 1 |
| ^1 (linear) | E | 0 | 47.5761668169349 | 123.000326114063 | 124.357766310499 | 126.041819332809 |
|  | T | 47.5761668169349 | 0 | 23.1750848855723 | 7.75466630858064 | 26.4736519020123 |
|  | M | 123.000326114063 | 23.1750848855723 | 0 | 18.876383586871 | 39.5077979627463 |
|  | D | 124.357766310499 | 7.75466630858064 | 18.876383586871 | 0 | 1 |
|  | C | 126.041819332809 | 26.4736519020123 | 39.5077979627463 | 1 | 0 |
| ^-2 (exp) | eu | 0 | 0.00044179531232 | 6.60978714623171E-05 | 6.46627506021843E-05 | 6.29463675606014E-05 |
|  | ta | 0.00044179531232 | 0 | 0.001861904205583 | 0.016629292454534 | 0.001426830196282 |
|  | mad | 6.60978714623171E-05 | 0.001861904205583 | 0 | 0.002806482974289 | 0.000640669946055 |
|  | dra | 6.46627506021843E-05 | 0.016629292454534 | 0.002806482974289 | 0 | 1 |
|  | cfr | 6.29463675606014E-05 | 0.001426830196282 | 0.000640669946055 | 1 | 0 |
| ^-1 (EXP) | eu | 0 | 0.021018927477866 | 0.00813005974531 | 0.008041315228381 | 0.00793387468773 |
|  | ta | 0.021018927477866 | 0 | 0.043149788013184 | 0.128954613932709 | 0.037773405939661 |
|  | mad | 0.00813005974531 | 0.043149788013184 | 0 | 0.052976249152699 | 0.025311458789552 |
|  | dra | 0.008041315228381 | 0.128954613932709 | 0.052976249152699 | 0 | 1 |
|  | cfr | 0.00793387468773 | 0.037773405939661 | 0.025311458789552 | 1 | 0 |
| ^-0.5 (EXP) | eu | 0 | 0.380761157100236 | 0.300277944470818 | 0.299455139918915 | 0.298449828847789 |
|  | ta | 0.380761157100236 | 0 | 0.455768878403725 | 0.599251626391339 | 0.440855777296096 |
|  | mad | 0.300277944470818 | 0.455768878403725 | 0 | 0.479755874732202 | 0.398868090381487 |
|  | dra | 0.299455139918915 | 0.599251626391339 | 0.479755874732202 | 0 | 1 |
|  | cfr | 0.298449828847789 | 0.440855777296096 | 0.398868090381487 | 1 | 0 |
| Niche similarity | E | 1 | 0.274 | 0.208 | 0.350 | 0.353 |
|  | T | 0.274 | 1 | 0.298 | 0.540 | 0.560 |
|  | M | 0.208 | 0.298 | 1 | 0.543 | 0.485 |
|  | D | 0.349 | 0.540 | 0.543 | 1 | 0.710 |
|  | C | 0.353 | 0.560 | 0.485 | 0.710 | 1 |

The standard BioGeoBEARS input script provided by Matzke (<http://phylo.wikidot.com/biogeobears#toc31>) was adjusted for the different models.

The script file for the Biogeographical Stochastic Mapping provided by Matzke (<http://phylo.wikidot.com/biogeographical-stochastic-mapping-example-script#toc12>) was adjusted to our best inferred model (the Drakensberg melting pot model with a manual dispersal multplier of 0.25 on the best tree).

The area adjacency matrix was set to the following:

|  | E | T | M | D | C |
| --- | --- | --- | --- | --- | --- |
| E | 1 | 1 | 0 | 0 | 0 |
| T | 1 | 1 | 1 | 1 | 0 |
| M | 0 | 1 | 1 | 1 | 0 |
| D | 0 | 1 | 1 | 1 | 1 |
| C | 0 | 0 | 0 | 1 | 1 |
