## Appendix 4 for "Leaps and bounds: geographical and ecological distance constrained the colonisation of the Afrotemperate by *Erica*": Appendix4.docx

**Appendix 4: Methods: Selected bootstrap trees used to represent phylogenetic uncertainty between geographically restricted major clades.**

1_o: (Cape-clade, *E. trimera*),(Dra-Ta-clade,(Ma-Ta-clade, *E. kingaensis*))

1_1: (Cape-clade, *E. trimera*),(Ma-Ta-clade,(Dra-Ta-clade, *E. kingaensis*))

1_2: (Cape-clade, *E. trimera*),(*E. kingaensis*,(Dra-Ta-clade,Ma-Ta-clade))

2_o: ((*E. trimera*,(Dra-Ta-clade,(Ma-Ta-clade, *E. kingaensis*))),Cape-clade)

2_1: ((*E. trimera*,(Ma-Ta-clade,(Dra-Ta-clade, *E. kingaensis*)),Cape-clade)

2_2: ((*E. trimera*,(*E. kingaensis*,(Dra-Ta-clade,Ma-Ta-clade)),Cape-clade)

o_o: ((Cape-Clade,(Dra-Ta-clade,(Ma-Ta-clade, *E. kingaensis*))), *E. trimera*)

o_1: ((Cape-Clade,(Ma-Ta-clade,(Dra-Ta-clade, *E. kingaensis*)), *E. trimera*)

o_2: ((Cape-Clade,(*E. kingaensis*,(Dra-Ta-clade,Ma-Ta-clade)), *E. trimera*)
