## Appendix 6 for "Leaps and bounds: geographical and ecological distance constrained the colonisation of the Afrotemperate by *Erica*"

**Appendix 6: Results: pairwise climate similarity (Schoener's *D*) between biogeographic areas.**

**Table S6.1.** Median of pairwise climate similarity (Schoener’s *D*) for PC **axis 1** between biogeographic areas.

| **Area** | **E** | **T** | **M** | **D** | **C** |
| --- | --- | --- | --- | --- | --- |
| **E** | 1 | 0.2084606 | 0.1713407 | 0.1869474 | 0.1889576 |
| **T** | 0.2084606 | 1 | 0.4258332 | 0.5315292 | 0.6437871 |
| **M** | 0.1713407 | 0.4258332 | 1 | 0.6761541 | 0.5410858 |
| **D** | 0.1869474 | 0.5315292 | 0.6761541 | 1 | 0.6031643 |
| **C** | 0.1889576 | 0.6437871 | 0.5410858 | 0.6031643 | 1 |

**Table S6.2.** Median of pairwise climate similarity (Schoener’s *D*) for PC **axis 2** between biogeographic areas.

| **Area** | **C** | **D** | **E** | **M** | **T** |
| --- | --- | --- | --- | --- | --- |
| **C** | 1 | 0.819248 | 0.5189766 | 0.4296022 | 0.4792726 |
| **D** | 0.819248 | 1 | 0.5180371 | 0.4133466 | 0.5508644 |
| **E** | 0.5189766 | 0.5180371 | 1 | 0.247694 | 0.3432868 |
| **M** | 0.4296022 | 0.4133466 | 0.247694 | 1 | 0.1723191 |
| **T** | 0.4792726 | 0.5508644 | 0.3432868 | 0.1723191 | 1 |
