## Appendix 7 for "Leaps and bounds: geographical and ecological distance constrained the colonisation of the Afrotemperate by *Erica*"

**Appendix 7: Results: Pairwise climate similarity (Schoener's *D*) between biogeographic areas for combined PC axes**

**Fig. S7.** Results of pairwise climate similarity (Schoener’s *D*) for combined PC axis 1 and 2. CFR, Cape; DRA, Drakensberg; EU, Europe; MAD, Madagascar; TA, Tropical Africa.

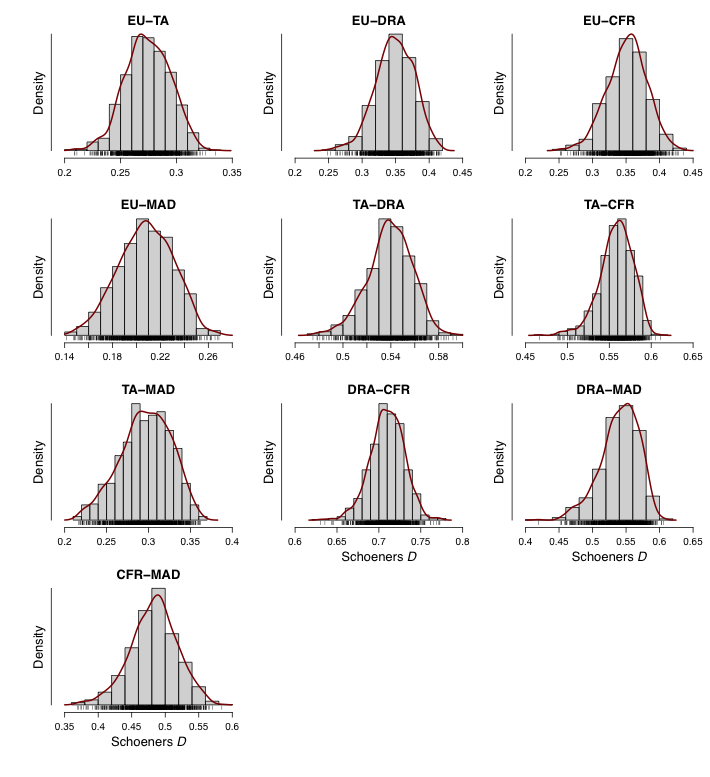

**Table S7.** Median of pairwise climate similarity (Schoener’s *D*) for combined PC axis 1 and 2 between biogeographic areas (used in the biogeographic analysis to define the niche similarity model).

| **Area** | **E** | **T** | **M** | **D** | **C** |
| --- | --- | --- | --- | --- | --- |
| **E** | 1 | 0.2741537 | 0.2082148 | 0.3488249 | 0.3527277 |
| **T** | 0.2741537 | 1 | 0.2981096 | 0.5396060 | 0.5599753 |
| **M** | 0.2082148 | 0.2981096 | 1 | 0.5434733 | 0.4846330 |
| **D** | 0.3488249 | 0.5396060 | 0.5434733 | 1 | 0.7104079 |
| **C** | 0.3527277 | 0.5599753 | 0.4846330 | 0.7104079 | 1 |
