## Appendix 8 for "Leaps and bounds: geographical and ecological distance constrained the colonisation of the Afrotemperate by *Erica*"

**Appendix 8:** Results of the different models: a) for the single best tree under DEC+J; b) for the single best tree under DEC; and c) for bootstrap trees under DEC+J. deltaAIC values are calculated overall across models for a given tree, and separately for the max areas/adjacency matrix, the biogeographic scenarios, and the distance models (“per comparison”). Models within deltaAIC=2 of the best score overall are bold underlined.

**a) Single best tree, DEC+J model.**

| Model | Dispersal multiplier | LnL | d [1/Ma] | e [1/Ma] | j | AIC | deltaAIC overall | deltaAIC per comparison |
| --- | --- | --- | --- | --- | --- | --- | --- | --- |
| no constraint |  | -69.98 | 0.0007 | 1E-12 | 0.0007 | 146 | 15 | 12.8 |
| Max area=2 |  | -69.9 | 0.0008 | 1E-12 | 0.0007 | 145.8 | 14.8 | 12.6 |
| Adjacency matrix |  | -63.61 | 0.0027 | 1E-12 | 0.0006 | 133.2 | 2.2 | 0 |
| Max area=2 + adjacency matrix |  | -63.6 | 0.0027 | 1E-12 | 0.0006 | 133.2 | 2.2 | 0 |
| The following models are based on the best model above | | | | | | | |  |
| Stepping Stone (w=1) | 0 | -96.18 | 0.008 | 1E-12 | 0.024 | 198.4 | 67.4 | 67.4 |
|  | 1 | -70.64 | 0.0054 | 1E-12 | 0.028 | 147.3 | 16.3 | 16.3 |
|  | 5 | -65.9 | 0.0059 | 1E-12 | 0.013 | 137.8 | 6.8 | 6.8 |
|  | 7.5 | -65.06 | 0.0061 | 1E-12 | 0.0085 | 136.1 | 5.1 | 5.1 |
|  | 10 | -64.52 | 0.0061 | 1E-12 | 0.0064 | 135 | 4 | 4 |
|  | **25** | **-63.34** | **0.0052** | **1E-12** | **0.0026** | **132.7** | **1.7** | **1.7** |
|  | **50** | **-63.14** | **0.004** | **1E-12** | **0.0013** | **132.3** | **1.3** | **1.3** |
| Cape to Cairo (w=1) | 0 | -83.07 | 0.0045 | 0.0006 | 0.0056 | 172.1 | 41.1 | 41.1 |
|  | 1 | -71.75 | 0.0035 | 1E-12 | 0.0032 | 149.5 | 18.5 | 18.5 |
|  | 5 | -67.49 | 0.0035 | 1E-12 | 0.0027 | 141 | 10 | 10 |
|  | 7.5 | -66.53 | 0.0035 | 1E-12 | 0.0025 | 139.1 | 8.1 | 8.1 |
|  | 10 | -65.89 | 0.0035 | 1E-12 | 0.0023 | 137.8 | 6.8 | 6.8 |
|  | 25 | -64.28 | 0.0034 | 1E-12 | 0.0017 | 134.6 | 3.6 | 3.6 |
|  | 50 | -63.63 | 0.0031 | 1E-12 | 0.0011 | 133.3 | 2.3 | 2.3 |
| Drakensberg Melting-pot (w=1) | 0 | -72.09 | 0.0033 | 0.0006 | 0.0037 | 150.2 | 19.2 | 19.2 |
|  | 1 | -64.09 | 0.0027 | 1E-12 | 0.0033 | 134.2 | 3.2 | 3.2 |
|  | **5** | **-62.52** | **0.0027** | **1E-12** | **0.0024** | **131** | **0** | **0** |
|  | **7.5** | **-62.62** | **0.0027** | **1E-12** | **0.0026** | **131.2** | **0.2** | **0.2** |
|  | **10** | **-62.52** | **0.0027** | **1E-12** | **0.0024** | **131** | **0** | **0** |
|  | **25** | **-62.51** | **0.0028** | **1E-12** | **0.0017** | **131** | **0** | **0** |

| (*E. aborea* European) | 25 | -61.92 | 0.0024 | 1E-12 | 0.0021 | 129.8 | x | x |
| --- | --- | --- | --- | --- | --- | --- | --- | --- |
| w =0.8 | **25** | **-62.61** | **0.0028** | **1E-12** | **0.0015** | **131.23** | **0.23** | **0.23** |
| w =0.5 | **25** | **-62.88** | **0.0028** | **1E-12** | **0.0011** | **131.76** | **0.76** | **0.76** |
| w =0.1 | **25** | **-63.43** | **0.0027** | **1E-12** | **0.00071** | **132.86** | **1.86** | **1.86** |
|  | **50** | **-62.88** | **0.0028** | **1E-12** | **0.0011** | **131.8** | **0.8** | **0.8** |
| Geographic distance | **As disp. probability (0 to 1)** | **-62.79** | **0.0029** | **1E-12** | **0.0012** | **131.58** | **0.58** | **0** |
|  | As distance (1 to x) | -72.37 | 0.0002 | 1E-12 | 0.00001 | 150.7 | 19.7 | 19.12 |
|  | As distance ^ -0.25 | -64.16 | 0.0034 | 1E-12 | 0.0013 | 134.3 | 3.3 | 2.72 |
|  | As distance ^ -1 | -71.40 | 0.004 | 1E-12 | 0.0032 | 148.8 | 17.8 | 17.22 |
|  | As distance ^ -2 | -83.18 | 0.0035 | 1E-12 | 0.0029 | 172.4 | 41.4 | 40.82 |
| Niche similarity | Schoener‘s D | -64.22 | 0.0047 | 1E-12 | 0.0013 | 134.4 | 3.4 | 2.82 |
| Niche plus distance as multipl. | As disp. probability/distance matrix | -64.01 | 0.0049 | 1E-12 | 0.0020 | 134.0 | 3.0 | 2.4 |

**b) Single best tree, DEC model.**

| Model | Dispersal multiplier | LnL | d [1/Ma] | e [1/Ma] | AIC | deltaAIC | deltaAIC per comparison |
| --- | --- | --- | --- | --- | --- | --- | --- |
| no constraint |  | -73.59 | 0.0011 | 1E-12 | 151.2 | 10 | 0.24 |
| Max area=2 |  | -73.48 | 0.0011 | 1E-12 | 150.96 | 9.76 | 0 |
| Adjacency matric |  | -76.72 | 0.0038 | 1E-12 | 157.44 | 16.24 | 6.48 |
| Max area=2 + adjacency matrix |  | -76.7 | 0.0039 | 1E-12 | 157.4 | 16.2 | 6.44 |
| The following models are based on the best model above | | | | | | |  |
| Stepping Stone (w=1) | 0 | -79.41 | 0.011 | 1E-12 | 162.8 | 21.6 | 21.6 |
|  | 1 | -76.63 | 0.01 | 1E-12 | 157.3 | 16.1 | 16.1 |
|  | 5 | -72.12 | 0.0076 | 1E-12 | 148.2 | 7 | 7 |
|  | 7.5 | -71.47 | 0.0066 | 1E-12 | 146.9 | 5.7 | 5.7 |
|  | 10 | -71.18 | 0.0058 | 1E-12 | 146.4 | 5.2 | 5.2 |
|  | 25 | -71.21 | 0.0034 | 1E-12 | 146.4 | 5.2 | 5.2 |
|  | 50 | -72.09 | 0.002 | 1E-12 | 148.2 | 7 | 7 |

| Cape to Cairo (w=1) | 0 | -89.65 | 0.0062 | 0.0005 | 183.3 | 42.1 | 42.1 |
| --- | --- | --- | --- | --- | --- | --- | --- |
|  | 1 | -76.7 | 0.0044 | 1E-12 | 157.4 | 16.2 | 16.2 |
|  | 5 | -73.14 | 0.0039 | 1E-12 | 150.3 | 9.1 | 9.1 |
|  | 7.5 | -72.43 | 0.0036 | 1E-12 | 148.9 | 7.7 | 7.7 |
|  | 10 | -72.03 | 0.0034 | 1E-12 | 148.1 | 6.9 | 6.9 |
|  | 25 | -71.52 | 0.0025 | 1E-12 | 147 | 5.8 | 5.8 |
|  | 50 | -72.09 | 0.0018 | 1E-12 | 148.2 | 7 | 7 |
| Drakensberg Melting-pot (w=1) | 0 | -77.02 | 0.004 | 0.0007 | 158 | 16.8 | 16.8 |
|  | 1 | -69.66 | 0.0035 | 1E-12 | 143.32 | 2.12 | 2.12 |
|  | **5** | **-68.66** | **0.0029** | **1E-12** | **141.3** | **0.1** | **0.1** |
|  | **7.5** | **-68.61** | **0.003** | **1E-12** | **141.2** | **0** | **0** |
| (*E. aborea* not widespread) | **7.5** | -68.54 | 0.003 | 1.00E-12 | 141.1 | x | x |
| w =0.8 | **7.5** | **-68.77** | **0.0028** | **1E-12** | **141.5** | **0.3** | **0.3** |
| w =0.5 | 7.5 | -69.67 | 0.0022 | 1E-12 | 143.3 | 2.1 | 2.1 |
| w =0.1 | 7.5 | -72.48 | 0.0013 | 1E-12 | 149 | 7.8 | 7.8 |
|  | **10** | **-68.66** | **0.0029** | **1E-12** | **141.3** | **0.1** | **0.1** |
|  | **25** | **-69.52** | **0.0023** | **1E-12** | **143** | **1.8** | **1.8** |
|  | 50 | -72.09 | 0.0018 | 1E-12 | 148.2 | 7 | 7 |
| Geographic distance | As disp multiplier (0 to 1) | -69.68 | 0.0021 | 1E-12 | 143.4 | 2.2 | 0 |
|  | As Distance ( to x) | -90.3 | 0.000018 | 1E-12 | 184.6 | 43.4 | 41.2 |
| (Divalike * J was better) | As distance ^ -0.25 | -71.38 | 0.0023 | 1E-12 | 146.8 | 5.6 | 3.4 |
|  | As distance ^ -1 | -74.66 | 0.0054 | 1E-12 | 153.3 | 12.1 | 9.9 |
|  | As distance ^ -2 | -85.62 | 0.0052 | 1E-12 | 175.2 | 34 | 31.8 |
| Niche similarity | Schoener‘s D | -72.84 | 0.0024 | 1E-12 | 149.67 | 8.47 | 6.27 |
| Niche plus distance as multipl. | As disp. probability/distance matrix | -70.42 | 0.0038 | 1E-12 | 144.8 | 3.6 | 1.4 |

**c) Bootstrap trees, DEC+J model.**

| BS-tree | Dispersal multiplier | Biogeographic model | LnL | AIC | deltaAIC | deltaAIC per comp. |
| --- | --- | --- | --- | --- | --- | --- |
| 1_1 |  | No constraint | -69.53 | 145.1 | 15.7 | 10.2 |
|  |  | Max area=2 + adjacency matrix | -64.43 | 134.9 | 5.5 | 0 |
|  | 0 | Drakensberg melting pot | -69.39 | 144.8 | 15.4 | 13.3 |
|  |  | Cape to Cairo | -73.68 | 153.4 | 24 | 21.9 |
|  |  | Stepping stone | -94.05 | 194.1 | 64.7 | 62.6 |
|  | 0.01 | Drakensberg melting pot | -65.69 | 137.4 | 8 | 5.9 |
|  |  | Cape to Cairo | -68.86 | 143.7 | 14.3 | 12.2 |
|  |  | Stepping stone | -75.05 | 156.1 | 26.7 | 24.6 |
|  | 0.1 | Drakensberg melting pot | -64.05 | 134.1 | 4.7 | 2.6 |
|  |  | Cape to Cairo | -63.51 | 133 | 3.6 | **1.5** |
|  |  | Stepping stone | -68.7 | 143.4 | 14 | 11.9 |
|  | 0.25 | Drakensberg melting pot | -63.89 | 133.8 | 4.4 | 2.3 |
|  |  | Cape to Cairo | -62.74 | 131.5 | 2.1 | **0** |
|  |  | Stepping stone | -67.03 | 140.1 | 10.7 | 8.6 |
|  | 0.5 | Drakensberg melting pot | -64.04 | 134.1 | 4.7 | 2.6 |
|  |  | Cape to Cairo | -63.13 | 132.3 | 2.9 | **0.8** |
|  |  | Stepping stone | -65.63 | 137.3 | 7.9 | 5.8 |
|  | distance based | **Niche similarity** | **-62.62** | **131.2** | **1.8** | **1.8** |
|  |  | Pure distance | -62.75 | 131.5 | 2.1 | 2.1 |
|  |  | **Niche similarity + pure distance** | **-61.71** | **129.4** | **0** | **0** |
| 1_2 |  | No constraint | -64.62 | 135.2 | 17.3 | 11.3 |
|  |  | Max area=2 + adjacency matrix | -58.97 | 123.9 | 6 | 0 |
|  | 0 | Drakensberg melting pot | -64.93 | 135.9 | 18 | 16.9 |
|  |  | Cape to Cairo | -67.88 | 141.8 | 23.9 | 22.8 |
|  |  | Stepping stone | -93.66 | 193.3 | 75.4 | 74.3 |
|  | 0.01 | Drakensberg melting pot | -59.16 | 124.3 | 6.4 | 5.3 |
|  |  | Cape to Cairo | -61.87 | 129.7 | 11.8 | 10.7 |
|  |  | Stepping stone | -69.51 | 145 | 27.1 | 26 |
|  | 0.1 | Drakensberg melting pot | -57.65 | 121.3 | 3.4 | 2.3 |
|  |  | **Cape to Cairo** | **-56.87** | **119.7** | **1.8** | **0.7** |
|  |  | Stepping stone | -62.61 | 131.2 | 13.3 | 12.2 |
|  | 0.25 | Drakensberg melting pot | -57.73 | 121.5 | 3.6 | 2.5 |
|  |  | **Cape to Cairo** | **-56.52** | **119** | **1.1** | **0** |
|  |  | Stepping stone | -60.91 | 127.8 | 9.9 | 8.8 |
|  | 0.5 | Drakensberg melting pot | -58.2 | 122.4 | 4.5 | 3.4 |
|  |  | Cape to Cairo | -57.3 | 120.6 | 2.7 | 1.6 |
|  |  | Stepping stone | -59.85 | 125.7 | 7.8 | 6.7 |
|  | distance based | Niche similarity | -57.19 | 120.4 | 2.5 | 2.5 |
|  |  | **Pure distance** | **-56.85** | **119.7** | **1.8** | **1.8** |
|  |  | **Niche similarity + pure distance** | **-55.93** | **117.9** | **0** | **0** |
| 1_o |  | No constraint | -68.72 | 143.4 | 19.9 | 11.4 |
|  |  | Max area=2 + adjacency matrix | -62.99 | 132 | 8.5 | 0 |
|  | 0 | Drakensberg melting pot | -76 | 158 | 34.5 | 34.5 |
|  |  | Cape to Cairo | -70.11 | 146.2 | 22.7 | 22.7 |
|  |  | Stepping stone | -99.88 | 205.8 | 82.3 | 82.3 |
|  | 0.01 | Drakensberg melting pot | -62.96 | 131.9 | 8.4 | 8.4 |
|  |  | Cape to Cairo | -62.14 | 130.3 | 6.8 | 6.8 |
|  |  | Stepping stone | -72.67 | 151.3 | 27.8 | 27.8 |
|  | 0.1 | Drakensberg melting pot | -61.51 | 129 | 5.5 | 5.5 |
|  |  | **Cape to Cairo** | **-58.73** | **123.5** | **0** | **0** |
|  |  | Stepping stone | -66.31 | 138.6 | 15.1 | 15.1 |
|  | 0.25 | Drakensberg melting pot | -61.65 | 129.3 | 5.8 | 5.8 |
|  |  | **Cape to Cairo** | **-59.45** | **124.9** | **1.4** | **1.4** |
|  |  | Stepping stone | -64.8 | 135.6 | 12.1 | 12.1 |
|  | 0.5 | Drakensberg melting pot | -62.17 | 130.3 | 6.8 | 6.8 |
|  |  | Cape to Cairo | -60.91 | 127.8 | 4.3 | 4.3 |
|  |  | Stepping stone | -63.85 | 133.7 | 10.2 | 10.2 |
|  | distance based | Niche similarity | -60.74 | 127.5 | 4 | 3.5 |
|  |  | Pure distance | -60.57 | 127.1 | 3.6 | 3.1 |
|  |  | **Niche similarity + pure distance** | **-59.02** | **124** | **0.5** | **0** |
| 2_1 |  | no constraint | -63.81 | 133.6 | 14.5 | 12.2 |
|  |  | no dispersal constraint | -57.69 | 121.4 | 2.3 | 0 |
|  | 0 | Drakensberg melting pot | -59.72 | 125.4 | 6.3 | 6.2 |
|  |  | Cape to Cairo | -66.2 | 138.4 | 19.3 | 19.2 |
|  |  | Stepping stone | -82.05 | 170.1 | 51 | 50.9 |
|  | 0.01 | Drakensberg melting pot | -58.06 | 122.1 | 3 | 2.9 |
|  |  | Cape to Cairo | -62.03 | 130.1 | 11 | 10.9 |
|  |  | Stepping stone | -67.05 | 140.1 | 21 | 20.9 |
|  | 0.1 | **Drakensberg melting pot** | **-56.59** | **119.2** | **0.1** | **0** |
|  |  | Cape to Cairo | -58.78 | 123.6 | 4.5 | 4.4 |
|  |  | Stepping stone | -59.95 | 125.9 | 6.8 | 6.7 |
|  | 0.25 | **Drakensberg melting pot** | **-56.61** | **119.2** | **0.1** | **0** |
|  |  | **Cape to Cairo** | **-57.5** | **121** | **1.9** | **1.8** |
|  |  | Stepping stone | -58.38 | 122.8 | 3.7 | 3.6 |
|  | 0.5 | **Drakensberg melting pot** | **-57** | **120** | **0.9** | **0.8** |
|  |  | **Cape to Cairo** | **-57.21** | **120.4** | **1.3** | **1.2** |
|  |  | Stepping stone | -57.74 | 121.5 | 2.4 | 2.3 |
|  | distance based | **Niche similarity** | **-57.57** | **121.1** | **2** | **2** |
|  |  | **Pure distance** | **-56.54** | **119.1** | **0** | **0** |
|  |  | **Niche similarity + pure distance** | **-57.18** | **120.4** | **1.3** | **1.3** |
| 2_2 |  | no constraint | x | 152.4 | 16.8 | 13.9 |
|  |  | no dispersal constraint | -66.27 | 138.5 | 2.9 | 0 |
|  | 0 | Drakensberg melting pot | -69.74 | 145.5 | 9.9 | 8.5 |
|  |  | Cape to Cairo | -72.49 | 151 | 15.4 | 14 |
|  |  | Stepping stone | -108.4 | 222.8 | 87.2 | 85.8 |
|  | 0.01 | Drakensberg melting pot | -67 | 140 | 4.4 | 3 |
|  |  | Cape to Cairo | -70.19 | 146.4 | 10.8 | 9.4 |
|  |  | Stepping stone | -74.48 | 155 | 19.4 | 18 |
|  | 0.1 | **Drakensberg melting pot** | **-65.54** | **137.1** | **1.5** | **0.1** |
|  |  | Cape to Cairo | -66.7 | 139.4 | 3.8 | 2.4 |
|  |  | Stepping stone | -68.75 | 143.5 | 7.9 | 6.5 |
|  | 0.25 | **Drakensberg melting pot** | **-65.51** | **137** | **1.4** | **0** |
|  |  | **Cape to Cairo** | **-65.57** | **137.1** | **1.5** | **0.1** |
|  |  | Stepping stone | -67.5 | 141 | 5.4 | 4 |
|  | 0.5 | **Drakensberg melting pot** | **-65.79** | **137.6** | **2** | **0.6** |
|  |  | **Cape to Cairo** | **-65.5** | **137** | **1.4** | **0** |
|  |  | Stepping stone | -66.66 | 139.3 | 3.7 | 2.3 |
|  | distance based | **Niche similarity** | **-65.75** | **137.5** | **1.9** | **1.9** |
|  |  | **Pure distance** | **-65.28** | **135.6** | **0** | **0** |
|  |  | **Niche similarity + pure distance** | **-65.18** | **136.4** | **0.8** | **0.8** |
| 2_o |  | no constraint |  | 148.6 | 18 | 13.5 |
|  |  | no dispersal constraint | -64.57 | 135.1 | 4.5 | 0 |
|  | 0 | Drakensberg melting pot | -66.06 | 138.1 | 7.5 | 6.9 |
|  |  | Cape to Cairo | -66.43 | 138.9 | 8.3 | 7.7 |
|  |  | Stepping stone | -96.87 | 199.7 | 69.1 | 68.5 |
|  | 0.01 | Drakensberg melting pot | -64.84 | 135.7 | 5.1 | 4.5 |
|  |  | Cape to Cairo | -64.22 | 134.4 | 3.8 | 3.2 |
|  |  | Stepping stone | -74.9 | 155.8 | 25.2 | 24.6 |
|  | 0.1 | Drakensberg melting pot | -63.66 | 133.3 | 2.7 | 2.1 |
|  |  | **Cape to Cairo** | **-62.88** | **131.8** | **1.2** | 0.6 |
|  |  | Stepping stone | -68.47 | 142.9 | 12.3 | 11.7 |
|  | 0.25 | Drakensberg melting pot | -63.71 | 133.4 | 2.8 | 2.2 |
|  |  | **Cape to Cairo** | **-62.59** | **131.2** | **0.6** | 0 |
|  |  | Stepping stone | -66.85 | 139.7 | 9.1 | 8.5 |
|  | 0.5 | Drakensberg melting pot | -64.04 | 134.1 | 3.5 | 2.9 |
|  |  | **Cape to Cairo** | **-63.14** | **132.3** | **1.7** | 1.1 |
|  |  | Stepping stone | -65.61 | 137.2 | 6.6 | 6 |
|  | distance based | **Niche similarity** | **-63.18** | **132.4** | **1.8** | **1.8** |
|  |  | **Pure distance** | **-62.76** | **131.5** | **0.9** | **0.9** |
|  |  | **Niche similarity + pure distance** | **-62.28** | **130.6** | **0** | **0** |
| o_1 |  | no constraint | -76.11 | 158.2 | 21.1 | 13.5 |
|  |  | no dispersal constraint | -69.33 | 144.7 | 7.6 | 0 |
|  | 0 | Drakensberg melting pot | -78.57 | 163.1 | 26 | 25.5 |
|  |  | Cape to Cairo | -78.26 | 162.5 | 25.4 | 24.9 |
|  |  | Stepping stone | -99.74 | 205.5 | 68.4 | 67.9 |
|  | 0.01 | Drakensberg melting pot | -69.74 | 145.5 | 8.4 | 7.9 |
|  |  | Cape to Cairo | -69.97 | 145.9 | 8.8 | 8.3 |
|  |  | Stepping stone | -77.47 | 160.9 | 23.8 | 23.3 |
|  | 0.1 | Drakensberg melting pot | -68.43 | 142.9 | 5.8 | 5.3 |
|  |  | **Cape to Cairo** | **-65.82** | **137.6** | **0.5** | 0 |
|  |  | Stepping stone | -72.32 | 150.6 | 13.5 | 13 |
|  | 0.25 | Drakensberg melting pot | -68.62 | 143.2 | 6.1 | 5.6 |
|  |  | **Cape to Cairo** | **-66.13** | **138.3** | **1.2** | **0.7** |
|  |  | Stepping stone | -71.46 | 148.9 | 11.8 | 11.3 |
|  | 0.5 | Drakensberg melting pot | -68.98 | 144 | 6.9 | 6.4 |
|  |  | Cape to Cairo | -67.39 | 140.8 | 3.7 | 3.2 |
|  |  | Stepping stone | -70.6 | 147.2 | 10.1 | 9.6 |
|  | distance based | Niche similarity | -66.94 | 139.9 | 2.8 | 2.8 |
|  |  | Pure distance | -67.19 | 140.4 | 3.3 | 3.3 |
|  |  | **Niche similarity + pure distance** | **-65.57** | **137.1** | **0** | **0** |
| o_2 |  | no constraint | -68.75 | 158.2 | 31.9 | 25.6 |
|  |  | no dispersal constraint | -63.31 | 132.6 | 6.3 | 0 |
|  | 0 | Drakensberg melting pot | -69.45 | 144.9 | 18.6 | 18.2 |
|  |  | Cape to Cairo | -70.82 | 147.6 | 21.3 | 20.9 |
|  |  | Stepping stone | -101.2 | 208.4 | 82.1 | 81.7 |
|  | 0.01 | Drakensberg melting pot | -62.61 | 131.2 | 4.9 | 4.5 |
|  |  | Cape to Cairo | -65.03 | 136.1 | 9.8 | 9.4 |
|  |  | Stepping stone | -73.64 | 153.3 | 27 | 26.6 |
|  | 0.1 | Drakensberg melting pot | -61.29 | 128.6 | 2.3 | 1.9 |
|  |  | **Cape to Cairo** | **-60.44** | **126.9** | **0.6** | **0.2** |
|  |  | Stepping stone | -66.54 | 139.1 | 12.8 | 12.4 |
|  | 0.25 | Drakensberg melting pot | -61.58 | 129.2 | 2.9 | 2.5 |
|  |  | **Cape to Cairo** | **-60.37** | **126.7** | **0.4** | **0** |
|  |  | Stepping stone | -64.91 | 135.8 | 9.5 | 9.1 |
|  | 0.5 | Drakensberg melting pot | -62.26 | 130.5 | 4.2 | 3.8 |
|  |  | Cape to Cairo | -61.4 | 128.8 | 2.5 | 2.1 |
|  |  | Stepping stone | -64 | 134 | 7.7 | 7.3 |
|  | distance based | Niche similarity | -61.61 | 129.2 | 2.9 | 2.9 |
|  |  | **Pure distance** | **-60.96** | **127.9** | **1.6** | **1.6** |
|  |  | **Niche similarity + pure distance** | **-60.17** | **126.3** | **0** | **0** |
| o_o |  | no constraint | -71.04 | 148.1 | 19.7 | 13.7 |
|  |  | no dispersal constraint | -64.2 | 134.4 | 6 | 0 |
|  | 0 | Drakensberg melting pot | -73.69 | 153.4 | 25 | 24 |
|  |  | Cape to Cairo | -72.9 | 151.8 | 23.4 | 22.4 |
|  |  | Stepping stone | -92.02 | 190 | 61.6 | 60.6 |
|  | 0.01 | Drakensberg melting pot | -64.96 | 135.9 | 7.5 | 6.5 |
|  |  | Cape to Cairo | -66.98 | 140 | 11.6 | 10.6 |
|  |  | Stepping stone | -73.31 | 152.6 | 24.2 | 23.2 |
|  | 0.1 | Drakensberg melting pot | -63.45 | 132.9 | 4.5 | 3.5 |
|  |  | **Cape to Cairo** | **-62.06** | **130.1** | **1.7** | **0.7** |
|  |  | Stepping stone | -67.8 | 141.6 | 13.2 | 12.2 |
|  | 0.25 | Drakensberg melting pot | -63.43 | 132.9 | 4.5 | 3.5 |
|  |  | **Cape to Cairo** | **-61.72** | **129.4** | **1** | **0** |
|  |  | Stepping stone | -66.57 | 139.1 | 10.7 | 9.7 |
|  | 0.5 | Drakensberg melting pot | -63.71 | 133.4 | 5 | 4 |
|  |  | Cape to Cairo | -62.51 | 131 | 2.6 | 1.6 |
|  |  | Stepping stone | -65.38 | 136.8 | 8.4 | 7.4 |
|  | distance based | Niche similarity | -62.33 | 130.7 | 2.3 | 2.3 |
|  |  | **Pure distance** | **-62.18** | **130.4** | **2** | **2** |
|  |  | **Niche similarity + pure distance** | **-61.18** | **128.4** | **0** | **0** |
