## Appendix 9 for "Leaps and bounds: geographical and ecological distance constrained the colonisation of the Afrotemperate by *Erica*"

**Appendix 9: Results: Summary of event counts from 50 biogeographical stochastic mappings under the best inferred model using the best tree.**

**a) DEC+J**

| **Mode** | **Type** | **Mean (SD)** | **%** |
| --- | --- | --- | --- |
| Within-area speciation | speciation | 473.2 (1.22) | 97.15 (0.25) |
|  | Speciation - subset | 3.6 (1.51) | 0.74 (0.31) |
| dispersal | Founder event (j) | 3.06 (0.84) | 0.63 (0.17) |
|  | Range expansion (d) | 5.12 (0.52) | 1.05 (0.11) |
|  | Range contraction (e) | 0 (0) | 0 (0) |
| Vicariance | vicariance | 2.12 (0.52) | 0.43 (0.11) |
| total |  | 487.1 (0.52) |  |

**b) DEC**

| **Mode** | **Type** | **Mean (SD)** | **%** |
| --- | --- | --- | --- |
| Within-area speciation | speciation | 471.3 (2.29) | 96.26 (0.47) |
|  | Speciation - subset | 6.06 (2.68) | 1.24 (0.55) |
| dispersal | Founder event (j) | 0 (0) | 0 (0) |
|  | Range expansion (d) | 7.64 (0.48) | 1.56 (0.1) |
|  | Range contraction (e) | 0 (0) | 0 (0) |
| Vicariance | vicariance | 4.64 (0.48) | 0.95 (0.47) |
| total |  | 489.6 (0.48) |  |
