## Appendix 10 for "Leaps and bounds: geographical and ecological distance constrained the colonisation of the Afrotemperate by *Erica*"

**Appendix 10: Number of range-expansion dispersal events (mean and standard deviation of all observed “d” dispersals)** **averaged across 50 biogeographical stochastic mappings under the best inferred model using the best tree. Rows represent the source area (where the lineages dispersed from) and columns the sink (where the lineages dispersed to). Abbreviations: E – Europe, T – Tropical Africa, M – Madagascar, D – Drakensberg, C – Cape.**

**a) DEC+J**

| **Area** | **E** | **T** | **M** | **D** | **C** | **Sum** | **%** |
| --- | --- | --- | --- | --- | --- | --- | --- |
| **E** | 0 (0) | 0.98 (0.14) | 0 (0) | 0 (0) | 0 (0) | 0.98 (0.14) | 19.14 (2.73) |
| **T** | 0.020 (0.14) | 0 (0) | 0.080 (0.27) | 0.98 (0.14) | 0 (0) | 1.08 (0.55) | 21.09 (10.74) |
| **M** | 0 (0) | 0 (0) | 0 (0) | 0 (0) | 0 (0) | 0 (0) | 0 (0) |
| **D** | 0 (0) | 0.16 (0.37) | 0 (0) | 0 (0) | 0 (0) | 0.16 (0.37) | 3.13 (7.23) |
| **C** | 0 (0) | 0 (0) | 0 (0) | 2.9 (0.30) | 0 (0) | 2.9 (0.3) | 56.64 (5.86) |
| **Sum** | 0.02 (0.14) | 1.14 (0.51) | 0.08 (0.27) | 3.88 (0.44) | 0 (0) | 5.12 (1.12) |  |
| **%** | 0.39 (2.73) | 22.27 (9.96) | 1.56 (5.27) | 75.78 (8.59) | 0 (0) |  | 100 (21.86) |

**b) DEC**

| **Area** | **E** | **T** | **M** | **D** | **C** | **Sum** | **%** |
| --- | --- | --- | --- | --- | --- | --- | --- |
| **E** | 0 (0) | 1 (0.2) | 0 (0) | 0 (0) | 0.02 (0.14) | 1.02 (0.34) | 13.35 (4.45) |
| **T** | 0.02 (0.14) | 0 (0) | 1 (0) | 1 (0) | 0 .98(0.14) | 3 (0.28) | 39.27 (3.66) |
| **M** | 0 (0) | 0 (0) | 0 (0) | 0 (0) | 0 (0) | 0 (0) | 0 (0) |
| **D** | 0 (0) | 0.6 (0.49) | 0 (0) | 0 (0) | 0 (0) | 0.6 (0.49) | 7.85 (6.41) |
| **C** | 0 (0) | 0.02 (0.14) | 0 (0) | 3(0) | 0 (0) | 3.02 (0.14) | 39.53 (.83) |
| **Sum** | 0.02 (0.14) | 1.62 (0.83) | 1 (0) | 4 (0) | 1 (0.28) | 7.64 (1.25) |  |
| **%** | 0.26 (1.83) | 21.20 (10.86) | 13.09 (0) | 52.36 (0) | 13.09 (22.4) |  | 100 (29.96) |
