## Appendix 11 for "Leaps and bounds: geographical and ecological distance constrained the colonisation of the Afrotemperate by *Erica*"

**Appendix 11:** **Number of cladogenetic dispersal events (mean and standard deviation of all observed jump 'j' dispersals) averaged from 50 biogeographical stochastic mappings under the best inferred model using the best tree. Rows represent the source area (where the lineage dispersed from) and columns the sink (where the lineage dispersed to). Abbreviations: E – Europe, T – Tropical Africa, M – Madagascar, D – Drakensberg, C – Cape.**

| **Area** | **E** | **T** | **M** | **D** | **C** | **Sum** | **%** |
| --- | --- | --- | --- | --- | --- | --- | --- |
| **E** | 0 | 0.040 (0.2) | 0 | 0 | 0 | 0.04 (0.2) | 1.31 (6.54) |
| **T** | 0 | 0 | 0.92 (0.27) | 0.040 (0.2) | 1 (0) | 1.96 (0.47) | 64.05 (15.36) |
| **M** | 0 | 0 | 0 | 0 | 0 | 0 | 0 |
| **D** | 0 | 0.82 (0.48) | 0 | 0 | 0 | 0.82 (0.48) | 26,780 (15.69) |
| **C** | 0 | 0.020 (0.14) | 0 | 0.22 (0.62) | 0 | 0.24 (0.76) | 7.84 (24.84) |
| **Sum** | 0 | 0.88 (0.82) | 0.92 (0.27) | 0.26 (0.82) | 1 (0) | 3.06 (1.24) |  |
| **%** | 0 | 28.76 (26,80) | 30.07 (8.82) | 8.50 (26.8) | 32.68 (0) |  | 100 |
