## Appendix 12 for "Leaps and bounds: geographical and ecological distance constrained the colonisation of the Afrotemperate by *Erica*"

**Appendix 12:** **Number of all dispersal events (mean and standard deviation of all observed anagenetic 'a' and 'd' dispersals, PLUS cladogenetic founder/jump dispersal) averaged from 50 biogeographical stochastic mappings under the best inferred model using the best tree. Note: anagenetic dispersal was always 0. Rows represent the source area (where the lineage dispersed from) and columns the sink (where the lineage dispersed to). Abbreviations: E – Europe, T – Tropical Africa, M – Madagascar, D – Drakensberg, C – Cape.**

**a) DEC + J**

| **Area** | **E** | **T** | **M** | **D** | **C** | **Sum** | **%** |
| --- | --- | --- | --- | --- | --- | --- | --- |
| **E** | 0 | 1.02 (0.14) | 0 | 0 | 0 | 1.02 (0.14) | 12.47 (1.71) |
| **T** | 0.02 (0.14) | 0 | 1 | 1.02 (0.14) | 1 | 3.04 (0.28) | 37.16 (3.42) |
| **M** | 0 | 0 | 0 | 0 | 0 | 0 (0) | 0 (0) |
| **D** | 0 | 0.98 (0.32) | 0 | 0 | 0 | 0.98 (0.32) | 11.98 (3.91) |
| **C** | 0 | 0.02 (0.14) | 0 | 3.12 (0.33) | 0 | 3.14 (0.47) | 38.39 (5.75) |
| **Sum** | 0.02 (0.14) | 2.02 (0.6) | 1 (0) | 4.14 (0.47) | 1 (0) | 8.18 (1.21) |  |
| **%** | 0.24 (1.71) | 24.69 (7.33) | 12.22 (0) | 50.61 (5.75) | 12.22 (0) |  | 100 (14.79) |

**b) DEC**

| **Area** | **E** | **T** | **M** | **D** | **C** | **Sum** | **%** |
| --- | --- | --- | --- | --- | --- | --- | --- |
| **E** | 0 (0) | 1 (0.2) | 0 (0) | 0 (0) | 0.02 (0.14) | 1.02 (0.34) | 13.35 (4.45) |
| **T** | 0.02 (0.14) | 0 (0) | 1 (0) | 1 (0) | 0 .98(0.14) | 3 (0.28) | 39.27 (3.66) |
| **M** | 0 (0) | 0 (0) | 0 (0) | 0 (0) | 0 (0) | 0 (0) | 0 (0) |
| **D** | 0 (0) | 0.6 (0.49) | 0 (0) | 0 (0) | 0 (0) | 0.6 (0.49) | 7.85 (6.41) |
| **C** | 0 (0) | 0.02 (0.14) | 0 (0) | 3(0) | 0 (0) | 3.02 (0.14) | 39.53 (.83) |
| **Sum** | 0.02 (0.14) | 1.62 (0.83) | 1 (0) | 4 (0) | 1 (0.28) | 7.64 (1.25) |  |
| **%** | 0.26 (1.83) | 21.20 (10.86) | 13.09 (0) | 52.36 (0) | 13.09 (22.4) |  | 100 (29.96) |
